# A first insight into the developability of an IgG3: A combined computational and experimental approach

**DOI:** 10.1101/2024.04.29.591602

**Authors:** Georgina B. Armstrong, Alan Lewis, Vidhi Shah, Paul Taylor, Craig J. Jamieson, Glenn A. Burley, Will Lewis, Zahra Rattray

## Abstract

Immunoglobulin G 3 (IgG3) monoclonal antibodies (mAbs) are high value scaffolds for developing novel therapies. Despite their wide-ranging therapeutic potential, IgG3 physicochemical properties and developability characteristics remain largely under-characterised. Protein-protein interactions elevate solution viscosity in high-concentration formulations impacting physico-chemical stability, manufacturability, and injectability of mAbs. Therefore, in this manuscript, the key molecular descriptors and biophysical properties of a model anti-IL-8 IgG1 and its IgG3 ortholog are characterised. A computational and experimental framework was applied to measure molecular descriptors impacting on their downstream developability. Findings from this approach underpin a detailed understanding of the molecular characteristics of IgG3 mAbs as potential therapeutic entities. This work is the first report examining the manufacturability of IgG3 for high concentration mAb formulations. While poorer conformational and colloidal stability, and elevated solution viscosity was observed for IgG3, future efforts controlling surface potential through sequence-engineering of solvent-accessible patches can be used to improve biophysical parameters that dictate mAb developability.

## INTRODUCTION

Antibody-based therapies possessing high specificity and superior efficacy have gained tremendous traction and growth in the biopharmaceuticals sector. Antibodies exert their pharmacological activity via a range of biological mechanisms-including and not limited to-direct blockade or activation of cell signal transduction pathways; Fc-mediated functions (antibody-dependent cell-mediated cytotoxicity,^1^ complement-dependent cytotoxicity, antibody-dependent cell phagocytosis); and immune activation.^2^ The molecular diversity of monoclonal antibody isotypes and subclasses can be harnessed to achieve different mechanisms of action in combating disease. Immunoglobulin G (IgG), the most abundant antibody isotype, can be further categorized as IgG1, IgG2, IgG3, and IgG4 subclasses in descending order of their prevalence in human serum.^3^

Whilst the sequence homology of IgG subclasses are highly conserved (>90%), each of these subclasses possess a unique hinge region length, differences in the number of inter-chain disulfide bonds and Fc-effector functionality.^3^ The molecular diversity of IgG subclasses and their involvement in mediating responses to different immunologic stimuli reflects the differing functional roles of IgG subclasses, affording their application in targeting a diverse antigen landscape. From a biotherapeutic perspective, there is growing recognition in recent years that the biomolecular properties of the different IgG subclasses correlate with improved developability characteristics, particularly in the context of targeting otherwise inaccessible biological targets.

Of the four IgG subclasses, IgG3 has the highest binding affinity for FcγRs, but is not routinely explored for therapeutic indications due to its historical suboptimal physicochemical stability profile and immunogenicity risk.^5^ However, the IgG3 hinge region influences the flexibility for this subclass of antibody, enabling IgG3 to interact more effectively with target antigens that are expressed at lower abundance.^6^ While both IgG1 and IgG3 play key roles in mediating immune responses, their structural differences lead to variations in their interactions with FcγRs and subsequent immune effector functions.^7^ IgG1 and IgG3 interact differently with most immune receptors (FcγR), triggering various immune effector mechanisms such as phagocytosis or antibody-dependent-cell-mediated cytotoxicity, which can offer therapeutic potential in immunooncology applications.^8^

IgG1 and IgG3 differ mostly based on the composition of their hinge region, which alters the extent of their ability to activate the immune system. IgG1 mAbs contain two inter-chain disulfide bonds in the hinge region, while IgG3 mAbs have 11 inter-chain disulfides. These structural differences influence their effector functions, with the IgG3 longer hinge length contributing to a combined greater accessibility to antigens and Fcγ receptors, resulting in more potent opsogenic activity.^5^

Beyond differences in their biological properties, each IgG subclass is associated with developability challenges, in the context of resistance to fragmentation, aggregation propensity, and elevated solution viscosity at high concentration.^4^ Though IgG1 mAbs exhibit superior stability under different pH conditions and in response to mechanical stress, they are more prone to fragmentation. However, IgG2 mAbs by comparison are less prone to fragmentation, but are more susceptible to aggregation.^4^

A dearth of IgG3 candidates in biopharmaceutical pipelines has been attributed to a lack of binding to protein A hampering downstream processing efforts,^9,10^ lack of *in vivo* stability resulting from proteolytic susceptibility, short plasma half-life necessitating a higher dosing frequency to achieve therapeutically-relevant levels,^11^ and immunogenicity concerns.^5,8^ However, with recent biotechnological advances in antibody sequence-based engineering, formulation strategies, and advancements in downstream processing these challenges can be mitigated. Mitigating such risks requires the development of IgG3-based molecular descriptors and biophysical properties under mAb formulation conditions, which identify key features which enhance as well as hinder downstream developability.

Here, a comprehensive study is presented to address the current knowledge gap of IgG3 developability characteristics, arising from sequence and structural differences to the IgG1 subclass. In this study, we analyse a computationally derived set of molecular descriptors of an anti-IL-8 IgG1 and IgG3 pair. This mAb pairing possesses identical variable domains. A comprehensive framework is then constructed to align the computational prediction of IgG1 and IgG3 sequences with measured experimental parameters evaluating their self-association behaviour and solution viscosity at high formulation concentration (>100 mg/mL).

## EXPERIMENTAL

### Computational methods

*In silico* homology modelling and antibody molecular descriptor calculations were performed in the Molecular Operating Environment (MOE) software, version 2020.0901 (Chemical Computing Group, Montreal, Canada).

### Homology modelling of anti-IL-8 IgG1 and IgG3

For both IgG1 and IgG3 molecules, full sequences of the heavy and light chains were inputted as the FASTA format into MOE (sequence editor) and annotated with a Kabat numbering scheme, with identical variable chain sequences. Constant chains were selected from the IMGT Repertoire database (https://www.imgt.org/IMGTrepertoire/), with accession numbers J00228 (IGHG1*01) and M12958 (IGHG3*01) for IgG1 and IgG3, respectively. For the IgG1 molecule, the Antibody modeller in MOE (version 2020.0901) was used to search for similar sequences with solved antibody structures to form the templates used for homology constructs. The variable fragment (Fv) of anti-IL-8 is published as PDB ID: 5OB5 (fAb complex with GroBeta). Fv fragments and full IgG structures were modelled by selecting ‘variable domain’ and ‘immunoglobulin’ model types, respectively. The immunoglobulin model type used the 1IGY PDB structure as a template to model the fragment crystallizable (Fc) region. A refinement gradient limit value of 1 was applied, and C-termini were capped with neutral residues, and superimposed to confirm structure alignment. For the IgG3, a different approach of independently modelling each antibody component was required due the absence of resolved IgG3 structures arising from the long hinge length. A new template hinge was generated independently using a mouse IgG2A (pdb:1IGT) as the second and fifth C-C disulfide bridges were in the same positions to the IgG3 hinge sequence (**supporting information**). This sequence was copied a further three times to generate four modules of the hinge. The Homology modeller in MOE (version 2020.0901) was used to generate 10 refined homology models for the hinge (**supporting information**). Each parameter was normalised to rank the geometric quality per model:

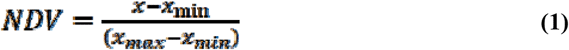

Where *NDV* is the normalised value for all geometric quality scores, except from the packing score, which was computed using equation 2.

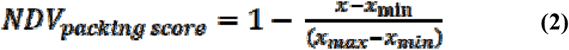

The lowest heavy atom root mean square deviation to the average position of intermediate models and lowest normalised score model were selected. A human Fc (pdb: 6D58) was imported for the Fc fragment, and the fragment antigen-binding regions (Fabs) were modelled *via* the Antibody modeller tool in MOE (version 2020.0901) from the anti-IL-8 IgG3 Fab sequence, with Fab selected as the model type. A 100% match to PDB ID 5OB5 was found as the variable sequence was the same between IgG1 and IgG3, with only a five-residue sequence difference in the constant regions of the Fab. All components were then joined manually and the joins energy minimised.

### Patch analysis of mAb1 IgG1 and IgG3 homology constructs

The protein patch tool in MOE was applied to each homology construct to identify electrostatic and hydrophobic surface patches. To aid visualisation of smaller surface patches, we set the following parameter thresholds: hydrophobic cut-off: ≥0.09 kcal/mol, hydrophobic min area: ≥30 Å^2^, charge cut-off: ≥30 kcal/mol/C, charge min area: ≥30 Å^2^, probe sphere radius: 1.8 Å.

### Predicted physicochemical descriptors

We computed a range of molecular descriptors (**supporting information**) for each full IgG1 and IgG3 model using the MOE *Protein Properties* tool. A NaCl concentration of 0.1 M was selected to represent the formulation buffer ionic strength at pH 6. *Hydrophobic imbalance* and *buried surface area* values were generated through BioMOE (version 2021-11-18, Chemical Computing Group, Montreal, Canada).

### Generation and biophysical analysis of mAb1 IgG1 and IgG3

#### IgG1 and IgG3 Expression and Downstream Purification

Chinese Hamster Ovary (CHO) K1 GS-KO (glutaminesynthetase-knockout) cells were used for to express IgG1 and IgG3. Heavy and light chain sequences were codon optimised and inserted into plasmids with CMV promoters by Atum Biosciences (Newark, CA,US). Plasmids were transfected *via* nucleofection with Leap-in Transposase® mRNA into Chinese Hamster Ovary (CHO) cells and maintained under selection conditions (no glutamine supplement) to generate stable pools. A fed-batch production process for 15 days with nutrient/glucose feeds every two or three days was deployed to increase expression of mAb1 IgG1 and IgG3. Cell culture bulks were fully clarified and then purified with an initial Protein L capture step followed by cation exchange polishing. Purified IgG1 and IgG3 were then concentrated, diafiltered and exchanged into formulation buffer containing histidine, trehalose, and arginine (pH 6) to a final target concentration of ≥150 mg/mL.

### Biophysical analysis of mAb1 IgG1 and IgG3

#### Analysis of identity

Peptide mapping was used to verify the full sequence identity for IgG1 and IgG3 (**supporting information**).

#### Analysis of purity

Analytical size-exclusion chromatography (aSEC) with UV-detection was deployed for monomeric purity assessment of mAb1 IgG1 and IgG3. A TSKgel Super SW3000, 4.6 x 300 mm (TOSOH Bioscience, United States) column was used with Agilent 1260 series HPLC (CA, US). Samples were prepared in water at 5 mg/mL and ran at 0.2 mL/min with a mobile phase containing 400 mM NaCl (pH 6.8). Chromatogram processing and integration was performed in The OpenLab CDS Data Analysis software (version 2.6, Agilent, California, US). The target monomeric purity of ≥95% was met by both mAb1 IgG1 and IgG3 molecules and aSEC was used to monitor physicochemical stability, by monitoring changes in the chromatogram.

#### Hydrophobic Interaction Chromatography (HIC) of IgG1 and IgG3

The hydrophobicity of IgG1 and IgG3 was assessed using HIC on an Agilent 1260 series HPLC (Agilent, California, US), coupled with UV detection (214 and 280 nm). A PolyLC PolyPROPUL 4.6 x 100 mm column was used, and to achieve separation based on net hydrophobicity, step-wise gradients of mobile phase B (low salt, with 50 mM ammonium sulfate) followed equilibration with mobile phase A (high salt,1.3 M ammonium sulfate). IgG1 and IgG3 samples were analysed at 1 mg/mL (5 μL injection volume) and a 0.7 mL/min flow rate.

#### Capillary Isoelectric Focusing (cIEF)

Charge distribution profiles of mAb1 IgG1 and IgG3 were assessed *via* capillary isoelectric focusing using an iCE3 instrument (Protein Simple, US). A range of pI markers (pI 3.85-8.77, Bio-Teche, Protein Simple, USA) were used to capture all acidic and basic isoforms for both molecules. To help prevent aggregation, 2M urea was added to the 1:1 ampholyte mixture (pH 3-10 and pH 8-10.5).The method entailed a pre-focus voltage of 1,500 V; an autosampler/transfer capillary temperature of 15 °C; a 10-12-minute focus voltage of 3,000 V; UV detection at 280 nm; a sample injection pressure of 2,000 mbar; a pre-focus time of 1 min; and a focus time of 10-12 minutes. The Empower 3 software (v4, Waters, US) was used for data analysis of peaks.

#### Zeta Potential

A Malvern Zetasizer (Malvern Panalytical, Malvern, UK) with a 633 nm laser was used to measure the zeta potential of the IgG1 and IgG3 pair by electrophoretic light scattering. Default settings included a 120 second equilibration time, automated attenuation and 10-100 measurement runs. There was a 60-second pause between each measurement and three technical replicate measurements were performed.

#### Determination of IgG1 and IgG3 Self-interaction

Self-association propensity of mAb1 IgG1 and IgG3 was measured with Affinity-Capture Self-Interaction Nanoparticle Spectroscopy (AC-SINS). Goat anti-human Fc and whole goat antibodies (Jackson ImmunoResearch, PA, USA) were prepared in 20 mM acetate buffer (pH 4.3), then mixed and incubated with 20 nm gold particles (Ted Pella Inc., CA, USA, concentration 7.0 x 10^11^ particles /mL). Test samples were prepared at 50 μg/mL in phosphate buffered saline (PBS) and 99 μL was added to 11 μL of nanoparticles in a 96 well plate, resulting in a final solution concentration of 50 μg/mL test mAb, 10x bead:anti-Fc conjugate and 0.02 mg/mL PEG2000. Plates were agitated, incubated for 2.5 hours and gently centrifuged to remove air bubbles. Absorbance measurements were read using a Pherastar FSX (BMG Labtech Ltd., Germany) plate reader, and spectra analysed with MARS software (v3.32, BMG Labtech Ltd., Germany). Differences in plasmon wavelengths for each sample were calculated from smoothed best fit curves. Experimental cutoffs included a <535 nm wavelength for negative controls (i.e., buffer).

#### Diffusion Self-interaction Parameter

A Stunner (Unchained Labs, CA, USA) dynamic light scattering setup was used to measure analyte hydrodynamic size, polydispersity, and diffusion coefficient. Data were analysed using the Lunatic & Stunner Client software (version 8.1.0.254). The measurement temperature was set as 25 □ with five, 10-second measurements acquired with a corresponding 1% extinction coefficient of 1.55AU*L/(g*cm) for all samples. Custom dispersant settings were applied (viscosity 1.26 cP and refractive index 1.33 at 20 °C) and both molecules were prepared in formulation buffer (0.5-20 mg/mL). The Lunatic & Stunner software (v8.1.0.244) were used for data export, and corresponding diffusion coefficients were used to calculate interaction parameters (k_D_) using linear regression plots.

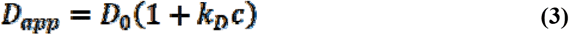

Where D_app_ refers to the apparent diffusion coefficient, D_0_ the self-diffusion coefficient at infinite dilution, and k_D_ the interaction parameter.

Exponential fits for diffusion coefficients over the test concentration range were used to calculate theoretical viscosities, adapted from the Generalised Stokes Einstein equation:

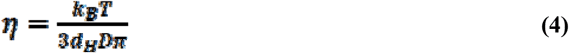

Where η is the theoretical dynamic viscosity (cP), *k*_*B*_*T* the Boltzmann constant at 298K, *d*_*H*_ the Z-ave diameter and *D* the diffusion coefficient.

#### Zeta Potential of mAb1 IgG1 and IgG3

A Malvern Zetasizer (Malvern Panalytical, Malvern, UK) with a 633 nm laser was used to measure the zeta potential of the IgG1 and IgG3 pair by electrophoretic light scattering. Each sample (refractive index 1.59) was prepared to 5 mg/mL in formulation buffer (pH 6.0, refractive index 1.33, viscosity at 1.26 cP) and a method was set up with equilibration time of 120 s, automatic attenuation and up to 100 runs per sample. A 60 s pause was also set between sample runs (a minimum of three technical replicates performed).

#### Analysis of Unfolding Temperatures

Differential scanning fluorimetry was performed on IgG1 and IgG3 mAb1 molecules using a Prometheus NT.48 setup (NanoTemper Technologies, Germany) with back-reflection technology. The intrinsic fluorescence from unfolding events exposing tyrosine and tryptophan residues were monitored *via* the 350/330 nm intensity ratio.^12^ A temperature ramp of 2°C/minute from 20-95 °C was performed. Both samples were assessed at concentrations ∼150 mg/mL and unfolding temperatures of antibody domains (T_m1_, T_m2_ and T_m3_) detected from first-derivative peaks of the 350/330 nm fluorescence intensity ratio. The first derivative peak of the scattering profile marked the aggregation temperature (T_agg_) values.

#### Measurement of Solution Viscosity

Viscosity curves were obtained using a VROC Initium (Rheosense, United States). The measurement protocol was optimised using the ‘Auto’ shear rate function, with fixed shear rates in the 100-2000 s^-1^ at each test concentration. Data were filtered to only include transient curves with steady plateaus with no drift and pressure over sensor position linear fits of R^2^ ≥0.998. Various models were used to fit the viscosity data. Firstly, the exponential-growth equation was applied:

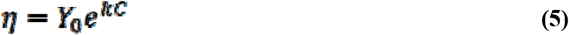

Where *η* is the dynamic viscosity (cP), *Y*_*0*_ the intercept, *k* the rate constant, and *c* the concentration of antibody (mg/mL).

Another model, developed by Tomar *et al*.^13,14^ was deployed for fitting the viscosity data:

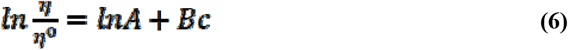

Where *η* is the dynamic viscosity (cP), *η*^*0*^ the buffer viscosity (cP) set at 1.13, *c* the concentration, and *lnA* the intercept of the slope *B*, when ln(η/η^0^) is plotted against concentration.

Finally, a modified Ross-Minton model was used to fit the viscosity-concentration profiles:

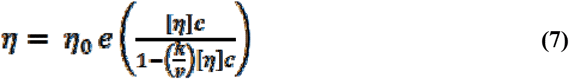

Where *η* is the dynamic viscosity (cP), *η*_0_ the buffer viscosity (cP) set at 1.13, *c* the concentration (mg/mL), [*η*] the intrinsic viscosity, *k* the crowding factor, *v* the Simha shape parameter. The[*η*], *k* and *v* parameters were estimated using the generalised reduced gradient (GRG) non-linear solver function to determine the local optimum reducing the sum of squared errors.

For intrinsic viscosity [*η*] measurements, multiple priming segments were set up followed by 10 replicates at the maximum shear rate of 23,080 s^-1^. Formulation buffer and anti-IL-8 formulations in the 5-50 mg/mL concentration range were measured to determine the relative viscosities (*η*_rel_) from which the specific (*η*_sp_) and reduced viscosities (*η*_red_) could be calculated (**supporting information**). The intrinsic viscosity was calculated from the linear regression of *η*_red_ over the sample concentration range tested, from which the Huggins coefficient was derived (Equation 7).

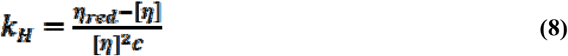

Where *k*_*H*_ is the Huggins Coefficient, *η*_red_ the reduced viscosity (cP) which is *η*_sp_/*c*, [*η*] the intrinsic viscosity (cP) and *c* the sample concentration (mg/mL).

The uncertainty of *k*_*H*_ was calculated from the propagation of error equation:

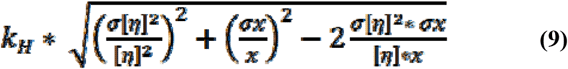

Where [*η*]^2^ the squared intrinsic viscosity, σ[*η*]^2^ the error of squared intrinsic viscosity, *x* the slope determined from the linear regression of *η*_red_ versus concentration, and σ*x* error of the slope.

#### Statistical Analysis

JMP Pro (v16.0.0, 2021) was used for multivariate analysis of computational predictions and measurement data to determine correlations between molecular descriptors and experimental parameters. We used GraphPad Prism (v5.04) for constructing graphs and performing unpaired t-test statistical analysis.

## RESULTS

### Patch analysis of homology constructs of anti-IL-8 IgG1 and IgG3

Solvent-accessible charge and hydrophobicity distribution profiles mAb self-association propensity that can promote aggregation.^15–17^ Disruption of hydrophobic patches has been previously correlated with reduced viscosity,^18,19^ driven by reduced native and non-native aggregation events.^20^ Furthermore, charge asymmetry between heavy and light chains has been correlated to increased self-association propensity, with increased electrostatic interactions.^15,21,22^ Therefore, we sought to assess the hydrophobic and electrostatic surface patch distribution profile for the anti-IL-8 IgG1 and IgG3 pair using full IgG homology constructs (**Figure 1**). Since the variable regions for both antibodies were similar, any differences occurring in the surface potential distributions were attributed to differences in the constant region between the molecules. Overall, both antibodies possessed a high proportion of hydrophobic patches (42% and 37%, respectively), with distinct differences in electrostatic patch (*i*.*e*., positive, and negative patch) distributions deriving predominantly from the increased residue exposure of the larger Fc domain of IgG3 (**supporting information**). The lowest energy conformation or the 62-residue IgG3 hinge region homology model was chosen (**supporting information**), contributing to 11% and 9% of the overall negative patch and positive residue contributions, respectively in comparison to the 4% and 1% contributions from the IgG1 hinge (**supporting information**).

**Figure 1.**
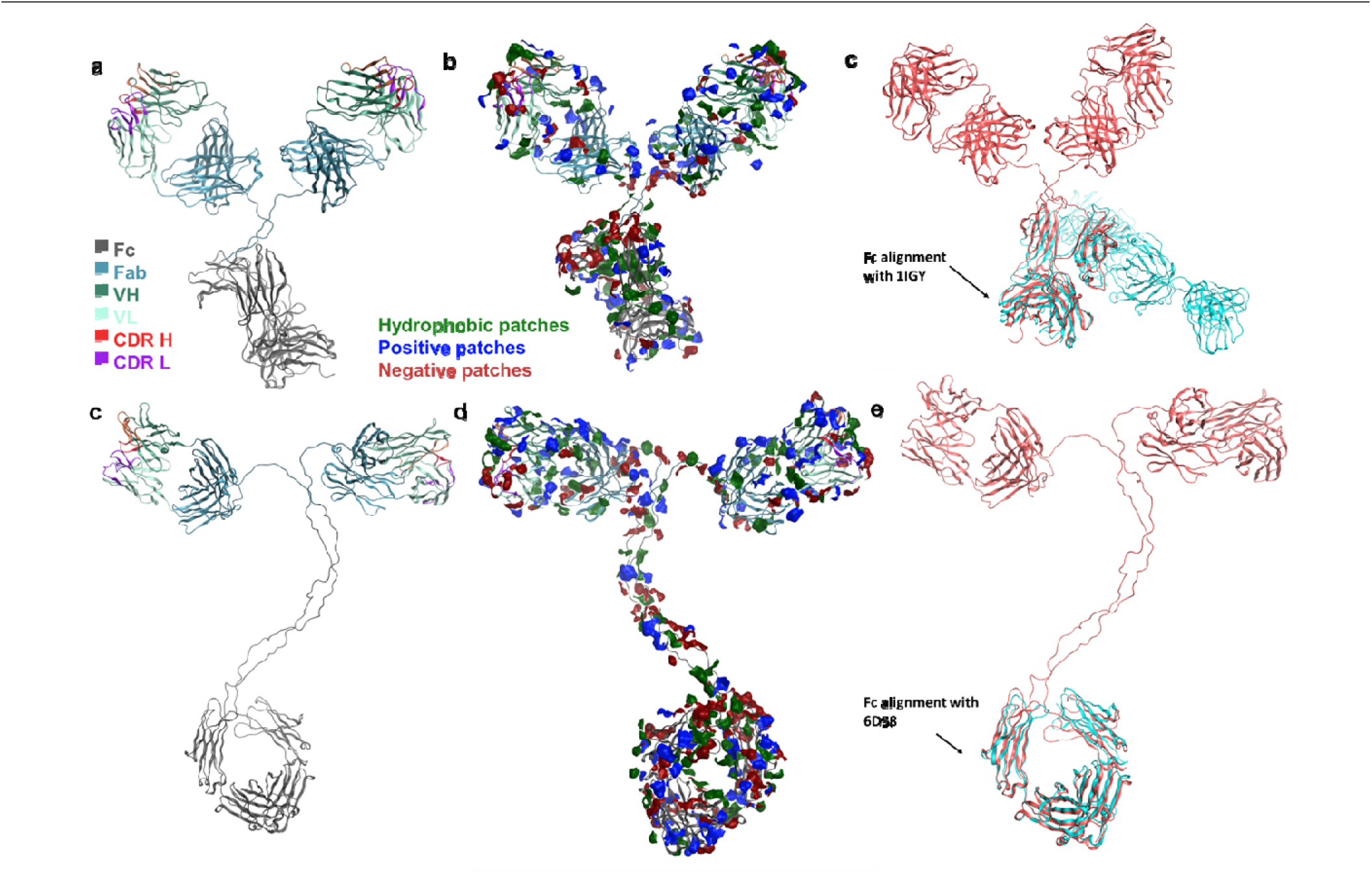
Homology constructs of the full IgG1 and IgG3 molecules. **a**, the full IgG1 structure **c**, the full IgG3 structure **b and d**, patch analysis of IgG1 and IgG3 homology constructs **c and e**, the Fc templates for IgG1 and IgG3.

### Biophysical analyses of mAb1 IgG1 and IgG3

#### Confirmation of identity and purity of mAb1 IgG1 and IgG3

To compare the biophysical properties of IgG3 to IgG1, a combined comprehensive pipeline consisting of computationally predicted molecular descriptors and experimental biophysical analyses was used We analysed the correlations between *in silico* and experimental charge, including hydrophobicity and colloidal parameters, and viscosity predictions and measurements. Both IgG1 and IgG3 sequence identities were confirmed with LC-MS peptide mapping (**supporting information**).

#### Antigen binding affinity of anti-IL-8 IgG1 and IgG3

The antigen affinity for the anti-IL-8 IgG1 and IgG3 antibody pair was assessed *via* surface plasmon resonance (SPR) (**Table 1**). Both molecules showed affinity (K_D_) for the IL-8 antigen with comparable association (k_a_) and dissociation (k_d_) rates. This demonstrated that the sequence and structural differences of the IgG3 constant domain had little influence on the Fv affinity for the target antigen.

**Table 1.** Antigen (IL-8) binding kinetics for IgG1 and IgG3 assessed *via* surface plasmon resonance (SPR). Corresponding (mean ± standard deviation) binding on-rate (k_a_), binding off-rate (k_d_) and the equilibrium dissociation constant (KD), the maximum response (R_max_) and goodness of fit (Chi-squared) of the 1:1 binding model. (N=3)

| Molecule | 1:1 binding kinetics | | | | Kinetics ($\chi^2$) |
| --- | --- | --- | --- | --- | --- |
| | $k_a \times 10^5 (M^{-1}s^{-1})$ | $k_d \times 10^{-4} (s^{-1})$ | $K_D (nM)$ | $R_{max} (RU)$ | $\chi^2$ |
| IgG1 | 3.84 ($\pm 0.12$) | 10.27 ($\pm 0.98$) | 2.67 ($\pm 0.16$) | 15.57 ($\pm 0.38$) | 1.57 ($\pm 0.62$) |
| IgG3 | 2.41 ($\pm 0.18$) | 9.17 ($\pm 0.05$) | 3.82 ($\pm 0.26$) | 14.63 ($\pm 0.15$) | 1.69 ($\pm 0.27$) |

#### Short term physical stability profiles of mAb1 IgG1 and IgG3

To be therapeutically viable, mAb formulations must have a solution phase stability of up to two years at refrigerated temperature and hours-several days under ambient storage conditions. A short-term stability study (up to 10 days) was conducted to assess relative changes in mAb1 monomeric purity from day 0 under refrigerated and ambient storage conditions (Figure 2). Both mAbs exhibited a reduction in monomeric purity from day 0 following storage temperatures from days 3-10. IgG3 showed a significant reduction in monomer purity from day 0 (surpassing the -2% drop threshold) when held at 25 °C by day 7, which could be attributed to increased soluble aggregate formation.

**Figure 2.**
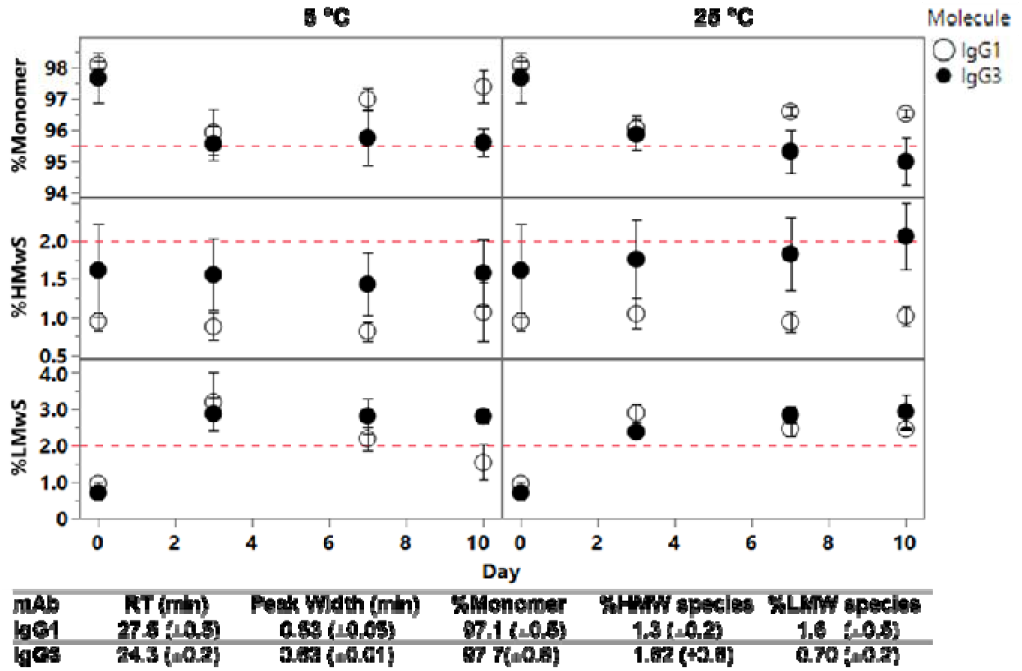
Reduced stability at 25 °C over 10 days for IgG3 compared to IgG1. Analytical size-exclusion chromatography was used to monitor the monomeric purity of mAb 1 IgG1 and IgG3 over 10 days at 5 °C and 25 °C. Delta differences relative to day 0 (%). Red dotted lines represent thresholds flagging changes in physical stability of mAbs. Corresponding monomeric purity and aggregate content as analysed by aSEC on day 0 for both molecules (bottom). Error bars represent standard deviations per sample, N=3.

Differential scanning fluorimetry (DSF) has been used as a surrogate for the assessment of mAb conformational stability and resistance to aggregation in previous work. Here, intrinsic fluorescence DSF was used to compare the unfolding temperatures IgG1 and IgG3 (**Figure 3**), with a lower temperature for the unfolding onset (T_onset_) and first unfolding event (T_m1_) being detected for IgG3, as well as a significant changes in the IgG3 thermal profile. No significant differences were detected for the temperature of aggregation onset (T_agg_), with IgG3 showing distinctly different scattering intensity profiles compared to IgG1 (**supporting information**). While there is a lack of published thermostability data on IgG3 molecules, the mAb1 IgG1 unfolding temperatures are in agreement with previous published values for IgG1 molecules in previous developability studies.^20^ The extended hinge region of IgG3 contributes to poor *in vivo* metabolic stability, increased number of allotypes and reduced half-life.^8,21,22^ Further studies on the conformational stability of mAb1 IgG3 are needed to develop more detailed insights for formulation shelf-life prediction in conjunction with functional stability and immunogenicity assessment. The immunogenicity of IgG3 resulting from its different glycoforms has been previously flagged for this subclass,^8^ necessitating monitoring of post-translational modifications over time.

**Figure 3.**
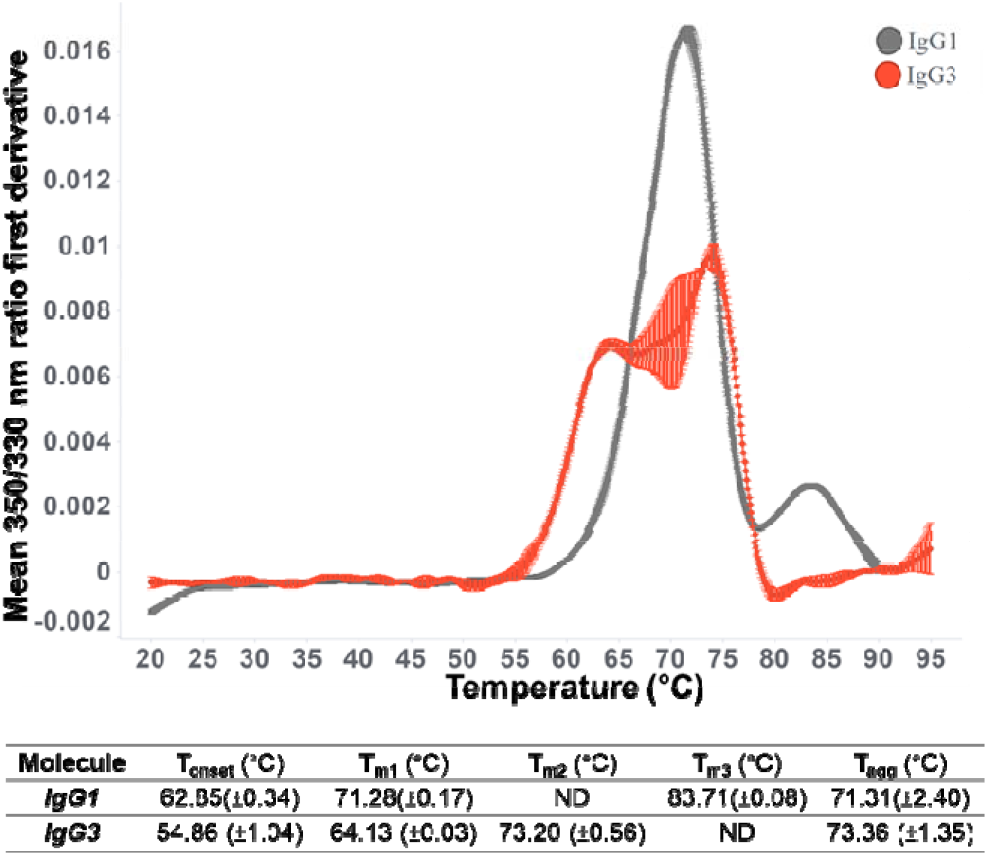
IgG3 shows reduced conformational stability compared to IgG1 at high concentrations. Thermal unfolding profiles for mAb1 IgG1 and IgG3. Mean first derivatives from the 350/330 nm ratio over a 20-95 °C temperature ramp reported as mean (±standard deviation). N=3.

#### Anti-IL-8 IgG3 has a positive charge under formulation conditions

Solvent accessible electrostatic patch distribution profiles of mAbs have previously been linked to changes in protein-protein interactions as a driver of self-association behaviour and elevated solution viscosity at high mAb formulation concentrations.^23,24^ We investigated how the predicted differences in electrostatic patch distribution profiles translated to measured charge parameters for anti-IL-8 IgG1 and IgG3 molecules (**Figure 4** and **Figure *5***). Comparable isoelectric points (pIs) (**Figure 4f**) were measured for IgG1 and IgG3; however, charge heterogeneity differences were observed with an increased proportion of acidic isoforms for IgG3 (**Figure 4c**), accompanied with an increased proportion of predicted negatively-charged patches in the constant domain. IgG1 and IgG3 showed significant differences in the mean measured zeta potential in formulation buffer at pH 6.0, with IgG3 had a positive zeta potential, whereas, IgG1 had a negative zeta potential (**Figure 4e**).

**Figure 4.**
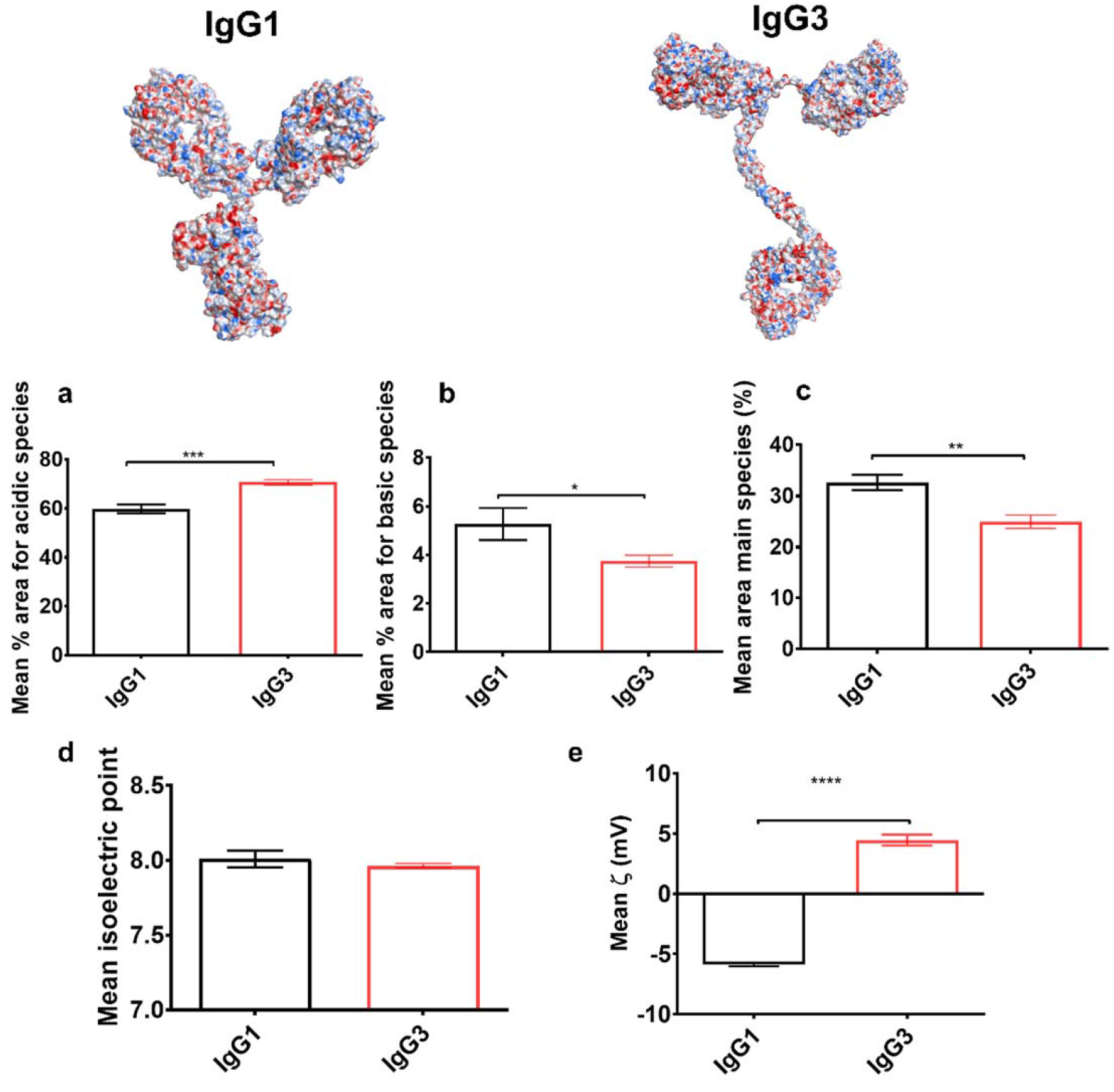
Different surface potential profiles were obtained for anti-IL-8 IgG1 and IgG3 predictions, which yielded comparable measured isoelectric points. Poisson-Boltzmann surface electrostatics were mapped onto homology constructs of anti-IL-8 IgG1 and IgG3, indicating regions of negative and positive charge density. Charge heterogeneity assessed *via* capillary isoelectric focussing (cIEF), **a**, acidic isoforms **b**, basic isoforms, **c**, main species, **d**, mean isoelectric point (pI), and **e**, zeta potential. Unpaired t-test **** denotes a P<0.0001, *** P<0.001, ** P<0.01, * P<0.1). Error bars represent standard deviation.

**Figure 5.**
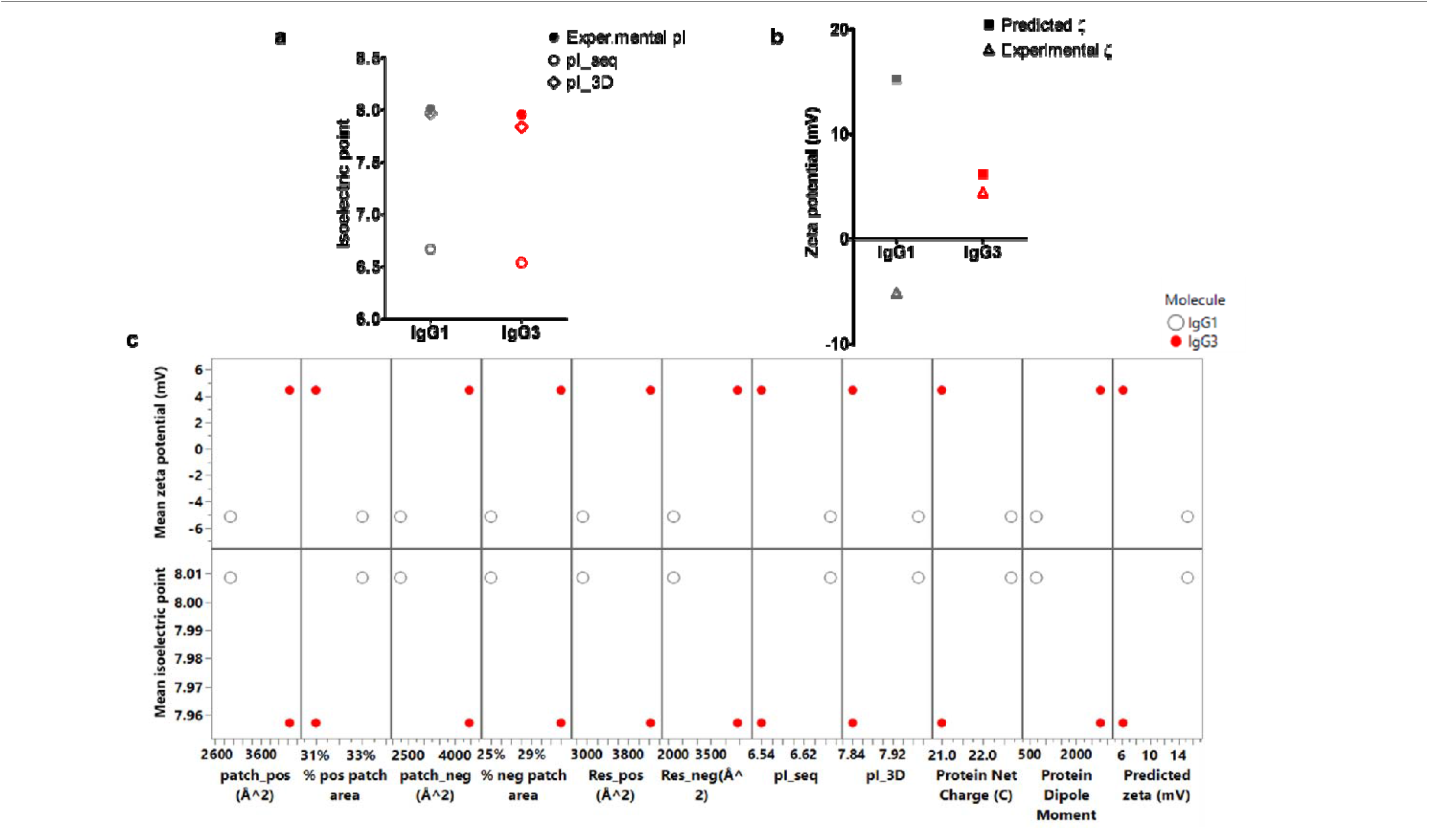
Correlating charge differences to *in silico* charge descriptors for anti-IL-8 IgG1 (grey) and IgG3 (red). The theoretical sequence-based pI was significantly lower than the experimentally measured pI. Zeta potential was measured at 5 mg/mL (**a**), demonstrating significant differences in electrical potential at the slipping plane between IgG1 (net negative charge) and IgG3 (net positive charge). Predicted zeta potential (computed at pH 6.0, 0.1 M NaCl) showed poor correlation with measured zeta potential values (**b**). Pair-wise comparisons between charge based in silico descriptors and experimental pI and zeta potential values was performed (**c**). Increased positive patch (patch_pos) and negative patch (patch_neg) areas, increased residue contributions to ionic patches (res_pos and res_neg), and decreased net charge aligned with experimental charge values.

The sequence and structure-based theoretical pIs predicted for IgG3 were slightly lower than those for IgG1, but the structure-based pI (pI_3D) directly correlated with experimental pI (**Figure *5*a**). Thorsteinson *et al*. similarly observed pI_3D to have the highest correlations to experimental parameters, but this was based off Fv models only and were also statistically comparable to the sequence-based pI method.^25^ Surprisingly, IgG3 showed a positive measured zeta potential (ζ) compared to the negative ζ for IgG1, contrary to predictions and isoelectric points (**Figure *5*b**). This suggests discrepancies between effective charge of the molecules in pH 6 formulation buffer and the net charge separated main species from capillary isoelectric point. The slight reduction observed in measured isoelectric point and increased measured zeta potential for IgG3 correlated with increased ionic patch area descriptors and reduced net charge (**Figure *5*c**). The increased negative patch count and area for IgG3 correlated with decreased predicted net charge which has been correlated previously with increased solution viscosity.^24,26,27^

#### Anti-IL-IgG3 exhibits a lower degree of hydrophobicity compared to IgG1

To evaluate the hydrophobicity of anti-IL-8 IgG1 and IgG3, hydrophobic interaction chromatography (HIC) was used (**Figure 6**). A significantly lower on-column retention time (RT) was observed for IgG3 in comparison to IgG1 (**Figure 6c**), aligning with most hydrophobic-based *in silico* descriptors and showing higher predicted hydrophobicity for IgG3 in comparison to IgG1 (with the exception of a slightly lower hydrophobic index and proportional percentage hydrophobic patch area) (**Figure 6e**). IgG3 also presented with increased peak broadening on the HIC column (**Figure 6d**), suggesting a potential increased population of different hydrophobic conformations. Increased net hydrophobicity has previously been correlated with increased solution viscosity occurring *via* cation-π and π-π stacking interactions from aromatic groups of solvent-exposed non-polar amino acid residues.^28,29^ Furthermore, increased hydrophobicity in the constant domain (Fc) of antibodies is widely correlated with a higher aggregation propensity, promoting an elevated mAb solution phase viscosity.^30,31^ Currently, there is a significant knowledge gap on drivers of IgG3 hydrophobicity, both measured and predicted, and how this affects the balance of domain-domain stability to unfolding propensity and aggregation.

**Figure 6.**
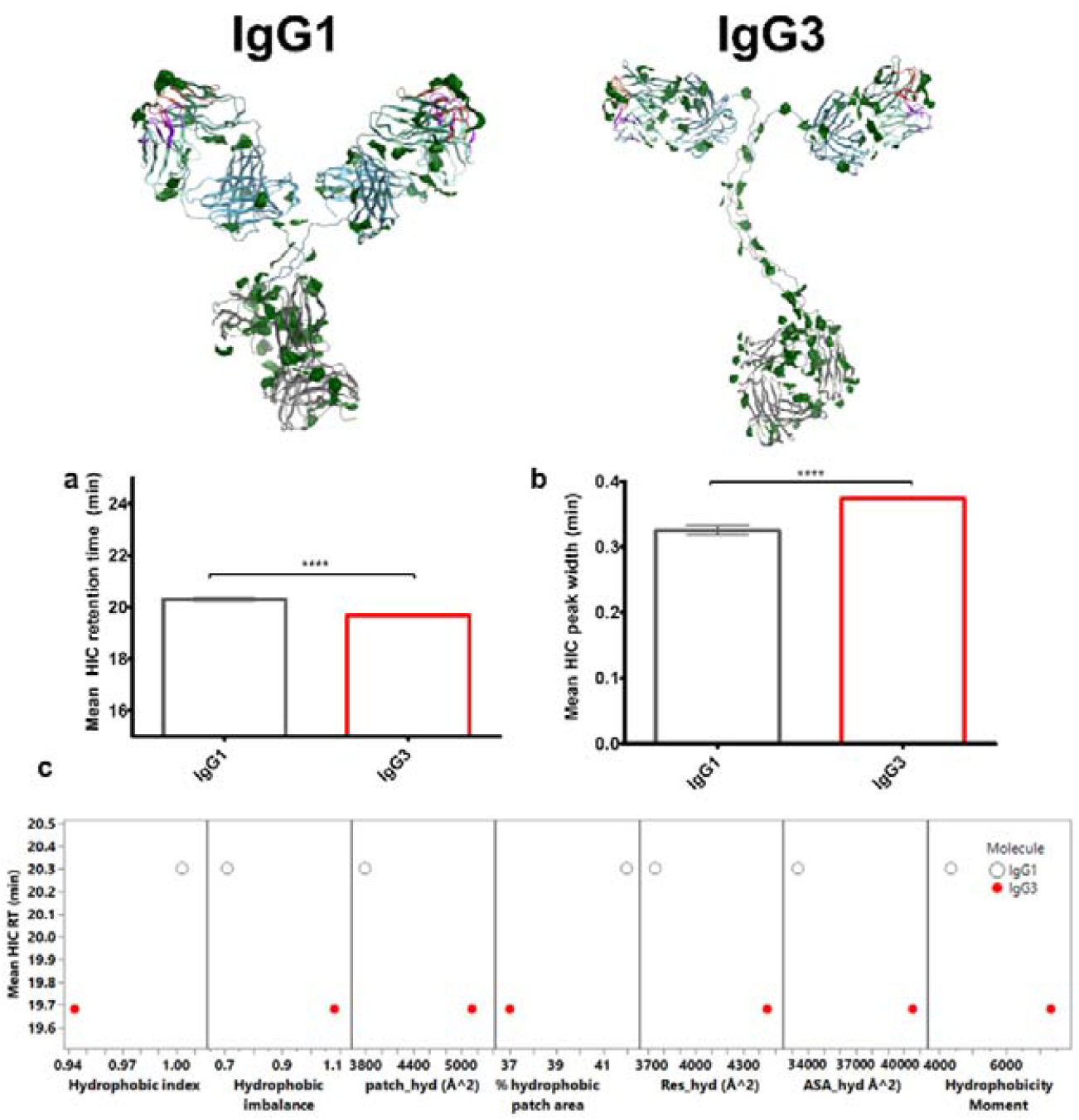
IgG3 exhibits a lower degree of hydrophobicity, contradicting computed solvent accessible hydrophobic area data. Protein patch surface maps for anti-IL-8 IgG1 and IgG3, filtered for hydrophobic patches (green). HIC chromatogram for **a**, IgG3 and **b**, IgG1. **c**, Pair-wise scatter plot comparisons between *in silico* descriptors and HIC retention time (RT). Unpaired t-test **** denotes a P<0.0001. Error bars represent standard deviation.

A significantly lower on-column retention time (RT) was observed for IgG3 in comparison to IgG1 (**Figure 6a**), disagreeing with most hydrophobic-based *in silico* descriptors that showed a higher predicted hydrophobicity for IgG3 compared to IgG1 (with the exception of the lower hydrophobic index and proportional percentage hydrophobic patch area) (**Figure 6c**). IgG3 also exhibited increased peak broadening on the HIC column (**Figure 6b**), which is indicative of a potential increase in hydrophobic conformations.

#### A comparison of anti-IL-8 IgG1 and IgG3 colloidal parameters

The concentration-dependent diffusion coefficient profile was measured for anti-IL-8 IgG1 and IgG3. We also used Affinity-Chromatography Self-Interaction Nanospectroscopy (AC-SINS) (**Figure 7**) as an orthogonal approach to measure the comparative self-association behaviour of IgG1 and IgG3. As expected, IgG3 measurements showed a larger hydrodynamic diameter (Z_ave) in comparison to IgG1, with a steady concentration-dependent increase over the 1-20 mg/mL test concentration range (**Figure 7a**), which corresponded to slower diffusion coefficients (**supporting information**). The measured self-interaction parameter, k_D_, for both molecules was negative and below the -15 mL/g threshold set, suggesting predominant attractive forces. However, the derived k_D_ parameter was significantly more negative for IgG3 anti-IL-8 compared to IgG1, indicative of increased self-association propensity. Conversely, IgG3 showed a comparable red shift from the AC-SINS assay, which did not correlate with the suggested increased self-association propensity from the DLS-derived k_D_ parameter.

**Figure 7.**
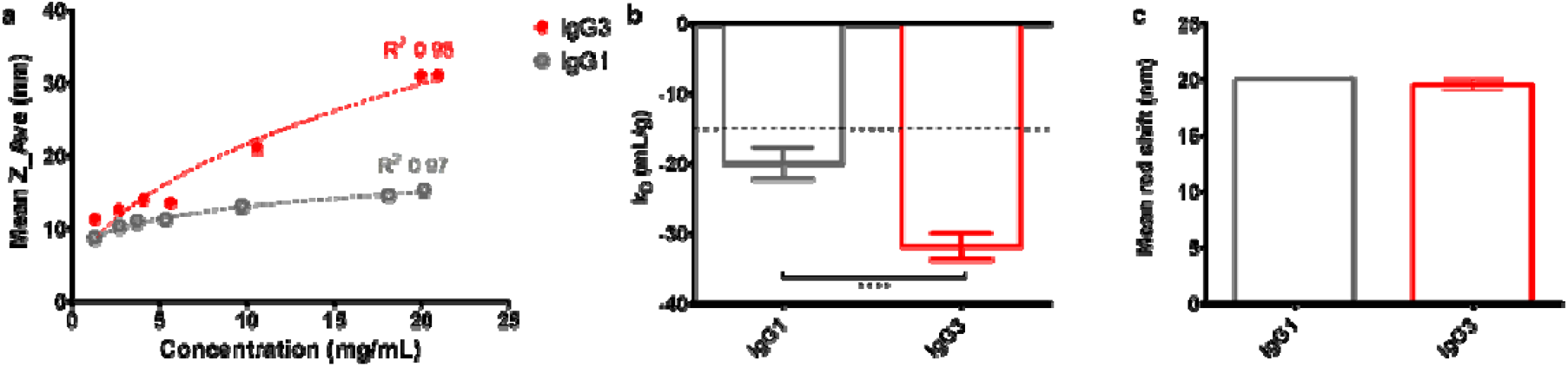
Colloidal interaction data from DLS measurements and AC-SINS for anti-IL-8 IgG1 and IgG3. **a**, concentration-dependent measured z-average hydrodynamic diameter. Logarithmic fits (10^(*Rope * log*(*X*)+*Y*)^) were applied to Z-Ave over concentration measurements, where Y was 0.92 and 0.87 and slopes were 0.2 and 0.46 for IgG1 and IgG3 respectively. Goodness of fit R^2^ values are reported. **b**, self-interaction parameter (k_D_) for IgG3. A dotted line at -15 mL/g represents a threshold for kD. **c**, Mean red shift in absorbance spectra from AC-SINS (N=2). Unpaired t-tests were performed to determine significant differences between means (**** denotes a P<0.0001). Error bars represent standard deviation. N=3.

#### Viscosity predictions and analysis

The Generalised Stokes Einstein viscosity (**Equation 4**) was calculated using DLS-derived diffusion coefficients (**supporting information**) and hydrodynamic diameters (**Figure 7a**). The resulting theoretical viscosities (**Figure 8**a) were log-transformed and showed a distinct increased viscosity for IgG3 at formulation concentrations ≥ 50 mg/mL in comparison to IgG1. Overestimation of the IgG3 viscosity and underestimation of IgG1 viscosity at 180 mg/mL (3,430 cP and 52 cP, respectively) is reflective of the derivation of data measured in the 1-20 mg/mL concentration regime, and the assumptions of using exponential fits for the diffusion coefficients and logarithmic fits for the Z-average values.

**Figure 8.**
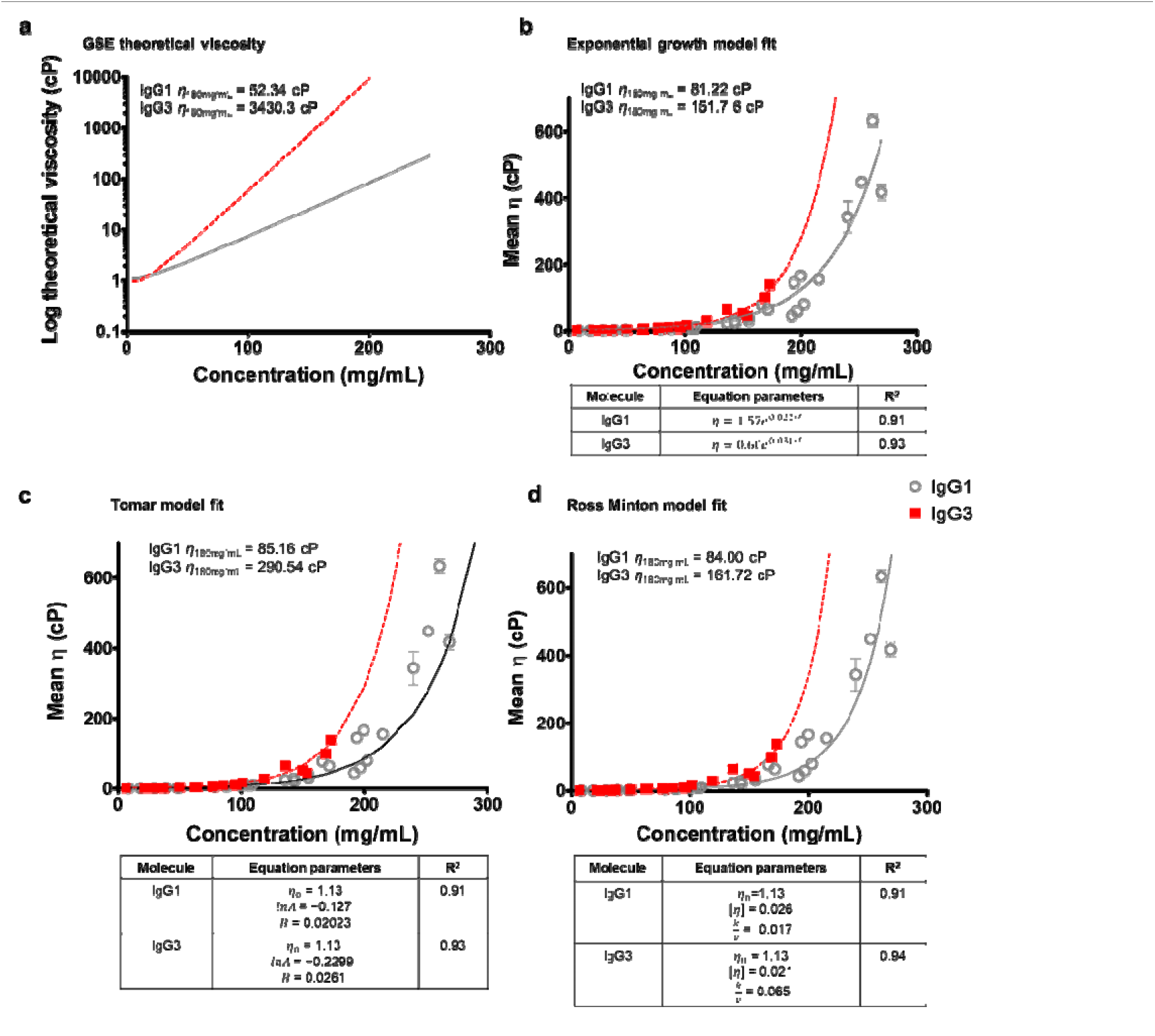
IgG3 has a higher apparent viscosity than IgG1 at high concentrations. **a**, the Generalised Stokes-Einstein equation was calculated from exponential extrapolation of diffusion coefficients and logarithmic fit of z-average diameters measured in the dilute range (1-20 mg/mL). Three viscosity model equations (lines) were used to fit the mean apparent viscosity data for IgG1 (grey circles) and IgG3 (black squares). **b**, the exponential growth model **c**, the modified Ross Minton model, and, **d**, the Tomar fit model. For each model, the predicted viscosity at 180 mg/mL is reported for both IgG1 and IgG3. Error bars represent standard deviation.

Therefore, we also measured apparent viscosities of IgG1 and IgG3 at concentrations up to 150 mg/mL (**Figure 8b-d**). We observed an elevated apparent viscosity for IgG3 compared to IgG1, in agreement with predicted theoretical viscosity and colloidal measurements. To compare the predictive power of different viscosity models, we fit the viscosity-concentration curves to three different models, including an exponential growth model (**Equation 5**), a Tomar model (Equation 6), and a modified Ross-Minton model (Equation 7). The exponential growth fit (**Figure 8b**) had a similar inflection point and gradient to the Ross-Minton fit (**Figure 8d**), resulting in similar viscosity interpolations at 180 mg/mL of 81.22 cP and 84 cP for IgG1, and 151.76 cP and 161.72 cP for IgG3, respectively. The Tomar model fit (**Figure 8c**) exhibited a shifted inflection point and steeper gradient compared to two previous models, resulting in higher interpolated viscosity predictions at 180 mg/mL (85.16 cP for IgG1 and 290.54 cP for IgG3).

Finally, we examined the individual contributions from each molecule to the solution viscosity by calculating intrinsic viscosity, [η], from measurements in the low concentration regime (0-50 mg/mL) (**Table 2**). Although statistically comparable to IgG1, IgG3 had an increased intrinsic viscosity, correlating with its increased hydrodynamic size. This suggests that the increased size and effective volume fraction of IgG3 increases the solution’s resistance to flow in the dilute regime.

**Table 2.** Intrinsic viscosity and Huggins coefficient for anti-IL-8 IgG1 and IgG3. Mean ± standard errors are shown. N=2.

| Molecule | Intrinsic viscosity (mL/g) | $k_H$ |
| --- | --- | --- |
| IgG1 | 8.28 (± 3.89) | 5.30 (± 1.77) |
| IgG3 | 10.42 (± 2.89) | 1.27(± 0.51) |

Moreover, the Huggins coefficient (k_H_) was computed, describing the changes in rate of viscosity increase from pairwise interactions. This has been previously equated to ‘solvent quality’ with values >0.5 suggestive of ‘poorer solvents’ that have solution viscosities more sensitive to protein-protein interactions (PPIs).^23^ Interestingly, IgG3 showed a reduction in k_H_ compared to IgG1, but both molecules had k_H_ >0.5, indicating poor solvation.^32^ Interestingly, IgG3 showed a reduction in k_H_ compared to IgG1, but both molecules had k_H_ >0.5, indicating poor solvation. It is worthwhile noting that the inaccuracies of the k_H_ parameter. Error in [η], from which k_H_ is derived, can arise from use of simple linear regression of η_red_/c (**supporting information**) as well as variability in viscosity measurements. Alternate non-linear fits may be able to account for antibody molecules which exceed the hard-sphere limit with regards to effective volume fraction of >2.5. kH has also been criticised to not account for solvation effects in dilute antibody solutions.^32–34^

## DISCUSSION

In this work we provide the first insights into the biophysical behaviour of a recombinant anti-IL8 IgG3, correlating *in silico* predicted molecular descriptors with experimental biophysical parameters and comparing these to a matched IgG1 with the same variable region sequence. Our goal was primarily to assess differences in physical stability and solution-phase viscosity-concentration profiles between these anti-IL-8 paired isotypes, while also predicting and measuring charge, hydrophobic and colloidal parameters as known drivers of mAb developability issues. Using a combined computational and experimental approach, we have constructed a set of guidelines that could be used more widely for mAb developability:

### (i) Reduced physical and conformational stability of anti-IL8 IgG3

We compared the short-term physical and thermal stability of IgG1 and IgG3 (Error! Reference source not found. and **Figure 3**), demonstrating a more rapid extent of monomer loss within a 10-day observation period. While there is a lack of published thermal stability data on IgG3, the IgG1 unfolding temperatures are broadly similar to published values for IgG1 molecules in prior developability studies.^24^ The extended hinge region of IgG3 is proposed to confer reduced *in vivo* stability, increased number of allotypes and reduced half-life.^5,25–27^ We hypothesise that the reduced domain unfolding temperatures we observed for IgG3 pair with the reported reduced conformational stability from the hinge region. Therefore, we propose additional structural analysis of anti-IL8 IgG3 conformational stability to better understand its role in formulation shelf-life prediction and pairing these findings with functional stability and immunogenicity assessment. The immunogenicity of IgG3 resulting from concerns on glycosylation propensity has previously been flagged for this subclass,^5^ necessitating the monitoring of IgG3 post-translational modifications over time between for both batch-to-batch and shelf-life stability.

### (ii) Predicted charge differences do not translate to differences in isoelectric points

Electrostatic surface potential mapping from homology constructs predicted an increase in the surface coverage of solvent accessible negatively-charged patches for anti-IL8 IgG3 in comparison to IgG1, suggesting an increased likelihood for electrostatic interactions to occur (**Figure 4** and **Figure *5*Error! Reference source not found**.). The theoretical isoelectric points (pIs) for IgG3 were predicted to be lower than IgG1. However, although slightly lower, the experimental pI for IgG3 was statistically comparable to IgG1. pI_3D showed a greater predictive power than pI_seq for the anti-IL8 full IgG models. Thorsteinson *et al*. similarly observed pI_3D to have the highest correlations to experimental parameters, but this was based on Fv models only and were statistically comparable to the sequence-based pI method.^28^ The increased negative patch count and area for IgG3 correlated with a decreased predicted net charge, which has been correlated previously with increased solution viscosity at dose-relevant formulation concentrations.^29–31^ Surprisingly, anti-IL8 IgG3 showed a positive measured zeta potential (ζ) compared to a negative potential for IgG1, which did not align with the *in silico* predictions of zeta potential and isoelectric points. This suggests discrepancies between the effective charge of the anti-IL8 IgG1 and IgG3 molecules in the pH 6 formulation buffer and the net charge separated main species from capillary isoelectric focusing.

### (iii) Net hydrophobicity of IgG3 does not correlate with predicted hydrophobic potential

Contrary to the predicted increased hydrophobic contributions from the hinge region, anti-IL8 IgG3 showed a shorter retention time on the hydrophobic-interaction chromatography column compared with IgG1 (**Figure 6**). Increased net hydrophobicity has previously been correlated with increased solution viscosity occurring *via* cation-π and π-π stacking interactions from aromatic groups of solvent-exposed non-polar amino acid residues.^19,32^ Moreover, increased hydrophobicity in the constant domain (Fc) of antibodies is widely correlated with a higher aggregation propensity, promoting an elevated mAb solution phase viscosity.^33,34^ In this case, as anti-IL8 IgG3 showed a decrease in net hydrophobicity, we cannot attribute the increased self-association or aggregation propensity to hydrophobic interactions. Currently, there is a significant knowledge gap on drivers of IgG3 hydrophobicity, both measured and predicted, and how this affects the balance of domain-domain stability to unfolding propensity and aggregation.

### (iv) Increased self-association propensity of anti-IL8 IgG3 correlates with hydrodynamic size and increased viscosity

The self-interaction parameter, k_D_, is widely used for predicting the propensity for protein-protein interactions at the molecular level, which drive elevated solution viscosity at high mAb formulation concentrations. For both molecules the k_D_ was negative and below the -15 mL/g arbitrary threshold set, suggesting predominant attractive forces. A more negative k_D_ was observed for anti-IL8 IgG3 (**Figure 7Error! Reference source not found**.), indicating more attractive interactions between molecules in the dilute concentration regime compared with IgG1.^35–38^

Unexpectedly, the AC-SINS red shift for IgG3, another metric used to experimentally predict mAb self-interaction propensity, showed a comparable absorbance intensity profile to the anti-IL8 IgG1. We hypothesise that increases in red shift may be masked by the reduced binding of IgG3 to the anti-Fc conjugated gold nanoparticles used during AC-SINS measurements. This may be a result of conformational flexibility provided by the extended IgG3 hinge region, leading to structural blocking of adjacent binding sites on the nanoparticles. Subsequently, this could reduce the number of bound antibodies to engage in self-interactions.

Across all viscosity fitting models, an increased apparent viscosity was observed for IgG3 in comparison to IgG1, aligning to the decreased predicted net charge, increased negative patch distributions, and increased hydrodynamic self-associations (**Figure 8**). The extrapolation of the Generalised Stokes-Einstein (GSE) model (**Figure 8a**) shows elevated viscosity, suggesting viscosity-contributing interactions in the dilute regime for anti-IL8 IgG3. This aligns to the increased intrinsic viscosity for IgG3 (**Table 2**), suggesting the increase hydrodynamic radius increases the fluid’s resistance to flow. Notably, no increase in the Huggin’s coefficient (k_H_) was observed for IgG3, which suggests comparable protein-protein pairwise interactions that contribute to IgG1 viscosity. However, it is worthwhile noting the inaccuracies of the k_H_ parameter. The error in [*η*], from which the k_H_ parameter is derived, can arise from the use of simple linear regression of *η*_red_/c fits (**supporting information**) as well as inter-experimental variability in viscosity measurements. Alternate non-linear fits may be able to account for antibody molecules, which exceed the hard-sphere limit with regards to effective volume fraction of >2.5. Another limitation of the Huggins coefficient is that it does not account for solvation effects in dilute antibody solutions. ^31,39,40^

It is important to note that our homology constructs are based on one possible conformation, and particularly with the assumed structure of IgG3, there are risks of under or overestimating the solvent-exposed surface potential. Our work uses these models as guiding tools to better understand mechanistic interactions that lead to molecular biophysical behaviour. There are growing efforts to research different structural modelling tools as well as use of molecular dynamics simulations with coarse grain simulation modelling^41,42^ that could help expand our knowledge of how both sequence and structure dictate interactions that lead to elevated viscosity and stability for IgG3.

## CONCLUSIONS

Pre-clinical developability assessment constitutes a prominent area of research for improving the probability of success for early-phase antibody candidates to reach clinical phases. Predictive tools probing the physicochemical and colloidal stability, affinity, and viscosity of antibodies in their formulation are being developed in combination with experimental assay pipelines as well as machine-learning algorithms. This work defines a multi-parameter set of guidelines for mAb using the context of biophysical behaviour of two IgG3 scaffolds as exemplars. We provide the first insights into the biophysical behaviour a recombinant anti-IL8 IgG3, comparing its computationally predicted molecular descriptors and experimentally-determined parameters to that of a paired IgG1 with the same variable region sequence. Our goal was primarily to assess the differences in physical stability and solution-phase viscosity-concentration profiles for these mAb1 paired isotypes as well as charge, hydrophobic and colloidal parameters. It is recognised that elevated solution viscosity of mAbs is driven by their self-association propensity. Hence, we used a combined *in silico* and comprehensive experimental pipeline to profile any viscosity differences between mAb1 IgG1 and IgG3 molecules. We reconciled the predicted computational descriptors derived from the *in silico* homology model, including the sequence and structure-based molecular descriptors determined for each mAb1 molecule, with their measured biophysical properties.

Here, we find that the constant domain of mAb1 IgG3 significantly influences its biophysical profile. IgG3 showed increased charge heterogeneity, hydrophobicity and self-association propensity, correlating with predicted increased hydrophobic and ionic surface potential from *in silico* homology modelling. This, alongside, decreased physical and conformational stability, aligns with the elevated solution viscosity observed for IgG3 compared with IgG1. The increased hydrodynamic size of IgG3 correlated with increased intrinsic viscosity, supporting increased thermodynamic as well as hydrodynamic contributions to solution viscosity.

Our work uniquely defines the bounds of manufacturability in the context of biophysical behaviour of an IgG3 molecule. We demonstrate the potential to further investigate the developability of the IgG3 subclass with formulation optimisations and/or *in silico* directed sequence-engineering. We propose future investigations are with functional assays to support the use of the IgG3 subclass as a promising therapeutic modality.

## ASSOCIATED CONTENT

### Supporting Information

The Supporting Information is available free of charge on the ACS Publications website.

Supporting information (PDF) contents:

Homology modelling of IgG3 (**Tables S1, S2**)

Patch analysis and physicochemical descriptor definitions (**Table S3**)

Patch analysis of anti-IL8 1 IgG1 and IgG3 (**Tables S4-5**) Physicochemical descriptors for anti-IL8 IgG1 and IgG3 (**Table S6**)

Analysis of identity by mass spectrometry (**Table S7**) Scattering intensity profiles from nano-differential scanning fluorimetry (nano-DSF) of anti-IL8 IgG1 and IgG3 (Figure S1)

Diffusion coefficients from dynamic light scattering (DLS) for anti-IL8 IgG1 and IgG3 (**Figure S2**)

Intrinsic viscosity and the Huggins coefficient for anti-IL8 IgG1 and IgG3 (**Figure S3**)

## AUTHOR INFORMATION

### Author Contributions

The manuscript was written through contributions of all authors. / All authors have given approval to the final version of the manuscript.

### Notes

GBA, AL, VS, PT and WL are employees of GlaxoSmithKline. The other authors declare no conflict of interest.

## ACKNOWLEDGMENTS

This study was sponsored by GlaxoSmithKline for Georgina Armstrong’s doctoral studies, the UK Engineering and Physical Sciences Research Council (EP/V028960/1), and the UK Biotechnology and Biological Sciences Research Council (BB/Y003268/1).

## REFERENCES

(1) Lo Nigro, C.; Macagno, M.; Sangiolo, D.; Bertolaccini, L.; Aglietta, M.; Merlano, M. C. NK-Mediated Antibody-Dependent Cell-Mediated Cytotoxicity in Solid Tumors: Biological Evidence and Clinical Perspectives. Ann. Transl. Med 2019, 7 (5), 105–105. 10.21037/atm.2019.01.42.

(2) Suzuki, M.; Kato, C.; Kato, A. Therapeutic Antibodies: Their Mechanisms of Action and the Pathological Findings They Induce in Toxicity Studies. J Toxicol Pathol 2015, 28 (3), 133–139. 10.1293/tox.2015-0031.

(3) Yu, J.; Song, Y.; Tian, W. How to Select IgG Subclasses in Developing Anti-Tumor Therapeutic Antibodies. J Hematol Oncol 2020, 13 (1), 45. 10.1186/s13045-020-00876-4.

(4) Cain, P.; Huang, L.; Tang, Y.; Anguiano, V.; Feng, Y. Impact of IgG Subclass on Monoclonal Antibody Developability. MAbs 2023, 15 (1), 2191302. 10.1080/19420862.2023.2191302.

(5) Chu, T. H.; Patz, E. F. J.; Ackerman, M. E. Coming Together at the Hinges: Therapeutic Prospects of IgG3. MAbs 2021, 13 (1), 1882028. 10.1080/19420862.2021.1882028.

(6) Plomp, R.; Dekkers, G.; Rombouts, Y.; Visser, R.; Koeleman, C. A. M.; Kammeijer, G. S. M.; Jansen, B. C.; Rispens, T.; Hensbergen, P. J.; Vidarsson, G.; Wuhrer, M. Hinge-Region O-Glycosylation of Human Immunoglobulin G3 (IgG3). Mol Cell Proteomics 2015, 14 (5), 1373–1384. 10.1074/mcp.M114.047381.

(7) Bolton, M. J.; Santos, J. J. S.; Arevalo, C. P.; Griesman, T.; Watson, M.; Li, S. H.; Bates, P.; Ramage, H.; Wilson, P. C.; Hensley, S. E. IgG3 Subclass Antibodies Recognize Antigenically Drifted Influenza Viruses and SARS-CoV-2 Variants through Efficient Bivalent Binding. Proc. Natl. Acad. Sci. U.S.A. 2023, 120 (35), e2216521120. 10.1073/pnas.2216521120.

(8) Damelang, T.; Rogerson, S. J.; Kent, S. J.; Chung, A. W. Role of IgG3 in Infectious Diseases. Trends in Immunology 2019, 40 (3), 197–211. 10.1016/j.it.2019.01.005.

(9) Boero, E.; Cruz, A. R.; Pansegrau, W.; Giovani, C.; Rooijakkers, S. H. M.; van Kessel, K. P. M.; van Strijp, J. A. G.; Bagnoli, F.; Manetti, A. G. O. Natural Human Immunity Against Staphylococcal Protein A Relies on Effector Functions Triggered by IgG3. Front Immunol 2022, 13, 834711. 10.3389/fimmu.2022.834711.

(10) Amaral, J.; Inganäs, M.; Cabral, J.; Prazeres, D. Study on the Scale-up of Human IgG3 Purification Using Protein A Affinity Chromatography. Bioseparation 2001, 10 (4), 139–143. 10.1023/A:1016353419499.

(11) Stapleton, N. M.; Andersen, J. T.; Stemerding, A. M.; Bjarnarson, S. P.; Verheul, R. C.; Gerritsen, J.; Zhao, Y.; Kleijer, M.; Sandlie, I.; de Haas, M.; Jonsdottir, I.; van der Schoot, C. E.; Vidarsson, G. Competition for FcRn-Mediated Transport Gives Rise to Short Half-Life of Human IgG3 and Offers Therapeutic Potential. Nat Commun 2011, 2, 599. 10.1038/ncomms1608.

(12) Kim, S. H.; Yoo, H. J.; Park, E. J.; Na, D. H. Nano Differential Scanning Fluorimetry-Based Thermal Stability Screening and Optimal Buffer Selection for Immunoglobulin G. Pharmaceuticals 2022, 15 (1), 29. 10.3390/ph15010029.

(13) Tomar, D. S.; Kumar, S.; Singh, S. K.; Goswami, S.; Li, L. Molecular Basis of High Viscosity in Concentrated Antibody Solutions: Strategies for High Concentration Drug Product Development. mAbs 2016, 8 (2), 216–228. 10.1080/19420862.2015.1128606.

(14) Tomar, D. S.; Li, L.; Broulidakis, M. P.; Luksha, N. G.; Burns, C. T.; Singh, S. K.; Kumar, S. In-Silico Prediction of Concentration-Dependent Viscosity Curves for Monoclonal Antibody Solutions. mAbs 2017, 9 (3), 476–489. 10.1080/19420862.2017.1285479.

(15) Yadav, S.; Laue, T. M.; Kalonia, D. S.; Singh, S. N.; Shire, S. J. The Influence of Charge Distribution on Self-Association and Viscosity Behavior of Monoclonal Antibody Solutions. Mol. Pharmaceutics 2012, 9 (4), 791–802. 10.1021/mp200566k.

(16) Nichols, P.; Li, L.; Kumar, S.; Buck, P. M.; Singh, S. K.; Goswami, S.; Balthazor, B.; Conley, T. R.; Sek, D.; Allen, M. J. Rational Design of Viscosity Reducing Mutants of a Monoclonal Antibody: Hydrophobic versus Electrostatic Inter-Molecular Interactions. mAbs 2015, 7 (1), 212–230. 10.4161/19420862.2014.985504.

(17) Dai, J.; Izadi, S.; Zarzar, J.; Wu, P.; Oh, A.; Carter, P. J. Variable Domain Mutational Analysis to Probe the Molecular Mechanisms of High Viscosity of an IgG1 Antibody. mAbs 2024, 16 (1), 2304282. 10.1080/19420862.2024.2304282.

(18) Geoghegan, J. C.; Fleming, R.; Damschroder, M.; Bishop, S. M.; Sathish, H. A.; Esfandiary, R. Mitigation of Reversible Self-Association and Viscosity in a Human IgG1 Monoclonal Antibody by Rational, Structure-Guided Fv Engineering. mAbs 2016, 8 (5), 941–950. 10.1080/19420862.2016.1171444.

(19) Tilegenova, C.; Izadi, S.; Yin, J.; Huang, C. S.; Wu, J.; Ellerman, D.; Hymowitz, S. G.; Walters, B.; Salisbury, C.; Carter, P. J. Dissecting the Molecular Basis of High Viscosity of Monospecific and Bispecific IgG Antibodies. MAbs 2020, 12 (1), 1692764. 10.1080/19420862.2019.1692764.

(20) Shan, L.; Mody, N.; Sormani, P.; Rosenthal, K. L.; Damschroder, M. M.; Esfandiary, R. Developability Assessment of Engineered Monoclonal Antibody Variants with a Complex Self-Association Behavior Using Complementary Analytical and in Silico Tools. Mol. Pharmaceutics 2018, 15 (12), 5697–5710. 10.1021/acs.molpharmaceut.8b00867.

(21) Raybould, M. I. J.; Marks, C.; Krawczyk, K.; Taddese, B.; Nowak, J.; Lewis, A. P.; Bujotzek, A.; Shi, J.; Deane, C. M. Five Computational Developability Guidelines for Therapeutic Antibody Profiling. Proceedings of the National Academy of Sciences 2019, 116 (10), 4025–4030. 10.1073/pnas.1810576116.

(22) Apgar, J. R.; Tam, A. S. P.; Sorm, R.; Moesta, S.; King, A. C.; Yang, H.; Kelleher, K.; Murphy, D.; D’Antona, A. M.; Yan, G.; Zhong, X.; Rodriguez, L.; Ma, W.; Ferguson, D. E.; Carven, G. J.; Bennett, E. M.; Lin, L. Modeling and Mitigation of High-Concentration Antibody Viscosity through Structure-Based Computer-Aided Protein Design. PLOS ONE 2020, 15 (5), e0232713. 10.1371/journal.pone.0232713.

(23) Yadav, S.; Shire, S. J.; Kalonia, D. S. Factors Affecting the Viscosity in High Concentration Solutions of Different Monoclonal Antibodies. JPharmSci 2010, 99 (12), 4812–4829. 10.1002/jps.22190.

(24) Bailly, M.; Mieczkowski, C.; Juan, V.; Metwally, E.; Tomazela, D.; Baker, J.; Uchida, M.; Kofman, E.; Raoufi, F.; Motlagh, S.; Yu, Y.; Park, J.; Raghava, S.; Welsh, J.; Rauscher, M.; Raghunathan, G.; Hsieh, M.; Chen, Y.-L.; Nguyen, H. T.; Nguyen, N.; Cipriano, D.; Fayadat-Dilman, L. Predicting Antibody Developability Profiles Through Early Stage Discovery Screening. MAbs 2020, 12 (1), 1743053. 10.1080/19420862.2020.1743053.

(25) Irani, V.; Guy, A. J.; Andrew, D.; Beeson, J. G.; Ramsland, P. A.; Richards, J. S. Molecular Properties of Human IgG Subclasses and Their Implications for Designing Therapeutic Monoclonal Antibodies against Infectious Diseases. Molecular Immunology 2015, 67 (2, Part A), 171–182. 10.1016/j.molimm.2015.03.255.

(26) Liu, H.; May, K. Disulfide Bond Structures of IgG Molecules: Structural Variations, Chemical Modifications and Possible Impacts to Stability and Biological Function. mAbs 2012, 4 (1), 17–23. 10.4161/mabs.4.1.18347.

(27) Li, W.; Prabakaran, P.; Chen, W.; Zhu, Z.; Feng, Y.; Dimitrov, D. S. Antibody Aggregation: Insights from Sequence and Structure. Antibodies 2016, 5 (3), 19. 10.3390/antib5030019.

(28) Thorsteinson, N.; Gunn, J. R.; Kelly, K.; Long, W.; Labute, P. Structure-Based Charge Calculations for Predicting Isoelectric Point, Viscosity, Clearance, and Profiling Antibody Therapeutics. MAbs 2021, 13 (1), 1981805. 10.1080/19420862.2021.1981805.

(29) Apgar, J. R.; Tam, A.; Sorm, R.; Moesta, S.; King, A.; Yang, H.; Kelleher, K.; Murphy, D.; D’Antona, A. M.; Yan, G.; Zhong, X.; Xiaotian Zhong; Xiaotian Zhong; Rodriguez, L.; Ma, W.; Ferguson, D.; Carven, G. J.; Gregory J. Carven; Bennett, E. M.; Eric M. Bennett; Laura Lin; Lin, L. Modeling and Mitigation of High-Concentration Antibody Viscosity through Structure-Based Computer-Aided Protein Design. PLOS ONE 2020, 15 (5), 1–26. 10.1371/journal.pone.0232713.

(30) Li, L.; Kumar, S.; Buck, P. M.; Burns, C.; Lavoie, J.; Singh, S. K.; Warne, N. W.; Nichols, P.; Luksha, N.; Boardman, D. Concentration Dependent Viscosity of Monoclonal Antibody Solutions: Explaining Experimental Behavior in Terms of Molecular Properties. Pharm Res 2014, 31 (11), 3161–3178. 10.1007/s11095-014-1409-0.

(31) Yadav, S.; Shire, S. J.; Kalonia, D. S. Factors Affecting the Viscosity in High Concentration Solutions of Different Monoclonal Antibodies. J Pharm Sci 2010, 99 (12), 4812–4829. 10.1002/jps.22190.

(32) Ausserwöger, H.; Schneider, M. M.; Herling, T. W.; Arosio, P.; Invernizzi, G.; Knowles, T. P. J.; Lorenzen, N. Non-Specificity as the Sticky Problem in Therapeutic Antibody Development. Nat Rev Chem 2022, 6 (12), 844–861. 10.1038/s41570-022-00438-x.

(33) Heads, J. T.; Kelm, S.; Tyson, K.; Lawson, A. D. G. A Computational Method for Predicting the Aggregation Propensity of IgG1 and IgG4(P) mAbs in Common Storage Buffers. mAbs 2022, 14 (1), 2138092. 10.1080/19420862.2022.2138092.

(34) Waibl, F.; Fernández-Quintero, M. L.; Wedl, F. S.; Kettenberger, H.; Georges, G.; Liedl, K. R. Comparison of Hydrophobicity Scales for Predicting Biophysical Properties of Antibodies. Front. Mol. Biosci. 2022, 9. 10.3389/fmolb.2022.960194.

(35) Chow, C.-K.; Allan, B. W.; Chai, Q.; Atwell, S.; Lu, J. Therapeutic Antibody Engineering To Improve Viscosity and Phase Separation Guided by Crystal Structure. Mol Pharm 2016, 13 (3), 915–923. 10.1021/acs.molpharmaceut.5b00817.

(36) Dynamic light scattering: a practical guide and applications in biomedical sciences - PMC. https://www.ncbi.nlm.nih.gov/pmc/articles/PMC5425802/ (accessed 2024-01-18).

(37) Lai, P.-K.; Austin Gallegos; Mody, N.; Sathish, H. A.; Trout, B. L. Machine Learning Prediction of Antibody Aggregation and Viscosity for High Concentration Formulation Development of Protein Therapeutics. mAbs 2022, 14 (1). 10.1080/19420862.2022.2026208.

(38) Dingfelder, F.; Henriksen, A.; Wahlund, P.-O.; Arosio, P.; Lorenzen, N. Measuring Self-Association of Antibody Lead Candidates with Dynamic Light Scattering (DLS). Therapeutic Antibodies: Methods and Protocols 2022, 241–258.

(39) Pathak, J. A.; Nugent, S.; Bender, M. F.; Roberts, C. J.; Curtis, R. J.; Douglas, J. F. Comparison of Huggins Coefficients and Osmotic Second Virial Coefficients of Buffered Solutions of Monoclonal Antibodies. Polymers (Basel) 2021, 13 (4), 601. 10.3390/polym13040601.

(40) Roche, A.; Gentiluomo, L.; Sibanda, N.; Roessner, D.; Friess, W.; Trainoff, S. P.; Curtis, R. Towards an Improved Prediction of Concentrated Antibody Solution Viscosity Using the Huggins Coefficient. Journal of Colloid and Interface Science 2022, 607, 1813–1824. 10.1016/j.jcis.2021.08.191.

(41) Prass, T. M.; Garidel, P.; Blech, M.; Schäfer, L. V. Viscosity Prediction of High-Concentration Antibody Solutions with Atomistic Simulations. J. Chem. Inf. Model. 2023, 63 (19), 6129–6140. 10.1021/acs.jcim.3c00947.

(42) Lai, P.-K.; Swan, J. W.; Trout, B. L. Calculation of Therapeutic Antibody Viscosity with Coarse-Grained Models, Hydrodynamic Calculations and Machine Learning-Based Parameters. mAbs 2021, 13 (1), 1907882. 10.1080/19420862.2021.1907882.

