## Supplemental for "A first insight into the developability of an IgG3: A combined computational and experimental approach"

1. Drug Substance Development, GlaxoSmithKline, Gunnels Wood Road, Stevenage, UK.
2. Computational and Modelling Sciences, GlaxoSmithKline, Gunnels Wood Road, Stevenage, UK
3. Large Molecule Discovery, GlaxoSmithKline, Gunnels Wood Road, Stevenage, UK
4. Department of Pure and Applied Chemistry, University of Strathclyde, Glasgow, UK.
5. Strathclyde Institute of Pharmacy and Biomedical Sciences, University of Strathclyde, Glasgow, UK.

#### Homology modelling of IgG3

**Table S1 IgG3 hinge was designed based off copied sequence of mouse IgG2A hinge (pdb 1IGT).** Residue sequence of 1IGT was used to aid construction of the IgG3 hinge. Cysteines are in highlighted in bold. The first 6 residues (yellow) contain two of the cysteines used to construct the IgG3 hinge in positions 2 and 5. To align the third cysteine for the anti-IL8 IgG3 hinge (position 11), the full 1IGT sequence was copied after the 6 residues (blue). This module was copied three more times to generate the four modules of the IgG3 hinge.

| | | | | | | | | | | | | | | | |
|---|---|---|---|---|---|---|---|---|---|---|---|---|---|---|---|
| 1IGT | 1 | **2** | 3 | 4 | **5** | 6 | | | | | | | 7 | 8 | 9 |
| | P | **C** | P | P | **C** | K | | | | | | | C | P | A |
| anti-IL8 IgG3 hinge module 1 | 1 | **2** | 3 | 4 | **5** | 6 | 7 | 8 | 9 | 10 | **11** | 12 | 13 | 14 | 15 |
| | P | **C** | P | P | **C** | K | P | C | P | P | **C** | K | C | P | A |

**Table S2 Hinge homology models for anti-IL8 IgG3.** 10 models were generated from the homology modeller tool in MOE (version 2020.0901) using the modified 1IGT hinge template (Table S1). Normalisation was performed for each quality parameter; normalisations to 1 used the minimum scores for RMSDs, minimum contact energy, highest packing score, minimum GB/VI interaction energies, minimum total potential energy (U), minimum solvation, electrostatic and van der Waal energies and minimum clashes and outliers. Model #2 was selected due to the lowest RMSD to Mean. This model also had the best (lowest) total normalised score. *RMSD to mean: heavy atom root mean square deviation to the average position of intermediate models, CA: alpha-carbon, GB/VI: Coulomb and Generalized Born interaction energies, U: total potential energy, E sol: solvation energy of the model, E ele: electrostatic energy of the model, E vdw: van der Waals energy of the model, BB; backbone*

| Model # | RMSD to Mean | CA RMSD to Mean | Contact Energy | Packing Score | GB/VI | U | E sol | E ele | E vdw | E bond | Atom Clashes | BB Bond Outliers | BB Angle Outliers | BB Torsion Outliers | Rotamer Outliers | Hinge model | Total Normalised Score |
|---|---|---|---|---|---|---|---|---|---|---|---|---|---|---|---|---|---|
| 1 | 0.63 | 0.47 | -21.97 | 1.98 | -4358.36 | -619.12 | -1458.85 | -2062.63 | -89.76 | 1533.27 | 93 | 10 | 21 | 20 | 1 | 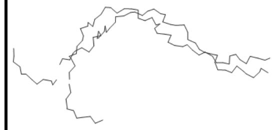 | 7.91 |
| 2 | 0.58 | 0.42 | -22.82 | 2.12 | -4047.83 | 686.84 | -2291.7 | -1042.57 | -36.16 | 1765.58 | 7 | 9 | 14 | 33 | 0 | 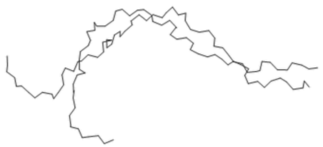 | 5.48 |
| 3 | 0.65 | 0.47 | -25.34 | 2.09 | -4039.06 | 731.05 | -2192.11 | -1018.76 | -14.76 | 1764.56 | 6 | 11 | 19 | 40 | 0 | 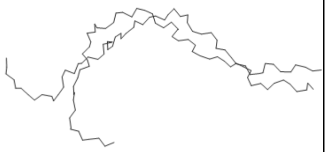 | 8.80 |
| 4 | 0.59 | 0.42 | -21.46 | 2.16 | -4036.56 | 688.62 | -2246.02 | -1036.27 | -36.85 | 1761.74 | 7 | 7 | 17 | 27 | 0 | 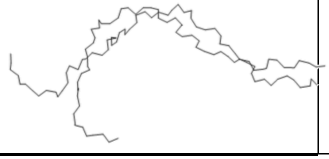 | 5.56 |

| | | | | | | | | | | | | | | | | | |
|---|---|---|---|---|---|---|---|---|---|---|---|---|---|---|---|---|---|
| 5 | 0.619 | 0.47 | -20.32 | 2.11 | -4033.33 | 805.83 | -2405.26 | -931.9 | -42.82 | 1780.55 | 7 | 11 | 16 | 29 | 0 | 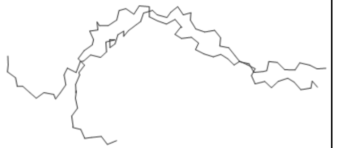 | 7.87 |
| 6 | 0.6 | 0.44 | -24.28 | 2.05 | -4032.27 | 796.83 | -2184.16 | -1008.91 | 1.019 | 1804.72 | 11 | 14 | 19 | 28 | 0 | 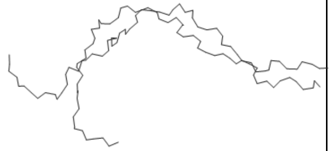 | 7.99 |
| 7 | 0.58 | 0.42 | -26.8 | 2.1 | -4004.76 | 1058.46 | -2466.78 | -864.31 | 29.79 | 1892.99 | 10 | 12 | 20 | 32 | 0 | 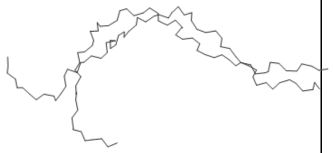 | 7.50 |
| 8 | 0.63 | 0.43 | -23.3 | 2.01 | -3993.05 | 871.17 | -2396.08 | -889.87 | -13.97 | 1775.01 | 9 | 10 | 14 | 33 | 0 | 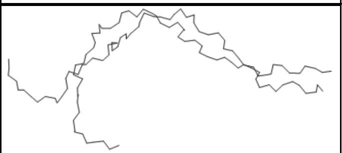 | 7.48 |
| 9 | 0.58 | 0.42 | -25.07 | 2.18 | -3977.17 | 1012.47 | -2545.39 | -802.08 | -14.35 | 1828.9 | 14 | 9 | 14 | 31 | 0 | 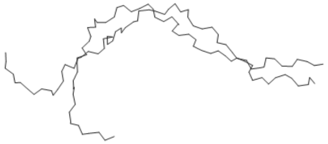 | 5.56 |
| 10 | 0.616 | 0.45 | -20.77 | 2.16 | -3967.21 | 1061.24 | -2467.2 | -815.9 | 58.66 | 1818.48 | 10 | 10 | 15 | 39 | 0 | 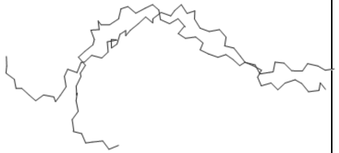 | 8.61 |

### Patch analysis and physicochemical descriptors

**Table S3** ***In silico*** **patch properties and physicochemical descriptors**

| Name | Description |
|---|---|
| **patch_hyd** | Summed area of hydrophobic patches (Å$^2$).[i] |
| **patch_hyd %** | Summed area of hydrophobic patches (Å$^2$) divided by summed area of all (ionic + hyd) patches (Å$^2$) |
| **patch_hyd_n** | Summed number of hydrophobic patches. [i] |
| **patch_pos Å2** | Summed area of positive patches (Å$^2$). **Error! Reference source not found.** |
| **patch_pos %** | Summed area of positive patches (Å$^2$) divided by summed area of all (ionic + hyd) patches (Å$^2$) |
| **patch_pos_n** | Summed number of positive patches. **Error! Reference source not found.** |
| **patch_neg Å2** | Summed area of negative patches (Å$^2$). **Error! Reference source not found.** |
| **patch_neg %** | Summed area of negative patches (Å$^2$) divided by summed area of all (ionic + hyd) patches (Å$^2$) |
| **patch_neg_n** | Summed number of negative patches. **Error! Reference source not found.** |
| **patch_ion** | Summed area of ionic (positive and negative) patches (Å$^2$). **Error! Reference source not found.** |
| **patch_ion_n** | Summed number of charged (positive and negative) patches. **Error! Reference source not found.** |
| **asa_hyd Å2** | Solvent-accessible surface area of hydrophobic atoms of a protein (Å$^2$). **Error! Reference source not found.** |
| **patch_cdr_hyd** | Summed area of hydrophobic patches near the CDRs (Å$^2$). **Error! Reference source not found.** |
| **patch_cdr_hyd_n** | Summed number of hydrophobic patches near the CDRs. **Error! Reference source not found.** |
| **patch_cdr_pos** | Summed area of positive patches near the CDRs (Å$^2$). **Error! Reference source not found.** |
| **patch_cdr_pos_n** | Summed number of positive patches near the CDRs. **Error! Reference source not found.** |
| **patch_cdr_neg** | Summed area of negative patches near the CDRs (Å$^2$). **Error! Reference source not found.** |
| **patch_cdr_neg_n** | Summed number of negative patches near the CDRs. [i] |
| **Hydrophobic Imbalance** | A vector that describes the displacement of the superficial geometric centre of the protein when the respective ASA values of each amino acid is considered.[ii] This was calculated through the descriptors function in the BioMOE module in MOE 2020. Default parameters were used with no sampling. This was calculated off the homology model at default values of pH 7.4, temperature of 300K and a salt concentration of 0.1M.[iii] |
| **Net_charge** | The formal protein net charge at a given pH. This was calculated through the Protein Properties tool in MOE 2020. The target pH was set to 6, temperature set to 300K and salt concentration to 0.1M.[i] |
| **Dipole_moment** | Dipole calculated across the protein from uneven distribution of charges. This was calculated through the Protein Properties tool in MOE 2020. The target pH was set to 6, temperature set to 300K and salt concentration to 0.1M. [i] |

| | |
|---|---|
| **Hyd_moment** | Hydrophobicity moment where each residue side chain hydrophobicity is calculated from the Kyte-Doolittle scale across the length of the protein.[iv] This was calculated through the Protein Properties tool in MOE 2020. The target pH was set to 6, temperature set to 300K and salt concentration to 0.1M.[i] |
| **Hydrophobicity Index** | The summation of hydrophobic residues' Eisenberg scores over the summation of hydrophilic residues' Eisenberg scores. Sharma et al. correlated Lower Eisenberg scores with lower viscosity.[v] |
| **Zeta potential** | Zeta potential is the electrical potential observed at the slipping plane. This was calculated through the Protein Properties tool in MOE 2020. The target pH was set to 6, temperature set to 300K and salt concentration to 0.1M.[i, vi] |
| **pI_seq** | The isoelectric point of a protein calculated from amino acid composition. This was calculated through the Protein Properties tool in MOE 2020. The target pH was set to 6, temperature set to 300K and salt concentration to 0.1M. [i,vii] |
| **pI_3D** | The isoelectric point of the molecule calculated through a modified version of Sillero's model. The PROPKA algorithm is used. This was calculated through the Protein Properties tool in MOE 2020. The target pH was set to 6, temperature set to 300K and salt concentration to 0.1M. [i] |
| **Res_ASA** | The summed contribution from each residue to the accessible surface area in Å$^2$. This was calculated through the Protein Properties tool in MOE 2020 and manually summed subsequently. The target pH was set to 6, temperature set to 300K and salt concentration to 0.1M. [i] |
| **Res_hyd** | The summed hydrophobic contribution from each residue to hydrophobic patch area in Å$^2$. This was calculated through the Protein Properties tool in MOE 2020 and manually summed subsequently. The target pH was set to 6, temperature set to 300K and salt concentration to 0.1M. [i] |
| **Res_pos** | The summed positive charge contribution from each residue to positive patch area in Å$^2$. This was calculated through the Protein Properties tool in MOE 2020 and manually summed subsequently. The target pH was set to 6, temperature set to 300K and salt concentration to 0.1M. [i] |
| **Res_neg** | The summed negative charge contribution from each residue to negative patch area in Å$^2$. This was calculated through the Protein Properties tool in MOE 2020 and manually summed subsequently. The target pH was set to 6, temperature set to 300K and salt concentration to 0.1M. [i] |
| **Hinge_res_ASA** | The summed contribution from each residue in the hinge region to the accessible surface area in Å$^2$. This was calculated from manually summing Res_ASA scores from residues in the hinge region. |
| **Hinge_res_hyd** | The summed hydrophobic contribution from each residue in the hinge to hydrophobic patch area in Å$^2$. This was calculated from |

| | |
|---|---|
| | manually summing Res_hyd scores from residues in the hinge region. |
| **Hinge_res_pos** | The summed positive charge contribution from each residue in the hinge to positive patch area in Å2. This was calculated from manually summing Res_pos scores from residues in the hinge region. |
| **Hinge_res_neg** | The summed negative charge contribution from each residue in the hinge to negative patch area in Å2. This was calculated from manually summing Res_neg scores from residues in the hinge region. |
| **Hinge_res_ASA contribution (%) to res_ASA** | $\frac{\text{Hinge\_res\_ASA}}{\text{Res\_ASA}} * 100$ |
| **Hinge_res_hyd contribution (%) to res_hyd** | $\frac{\text{Hinge\_res\_hyd}}{\text{Res\_hyd}} * 100$ |
| **Hinge_res_pos contribution (%) to res_pos** | $\frac{\text{Hinge\_res\_pos}}{\text{Res\_pos}} * 100$ |
| **Hinge_res_neg contribution (%) to res_neg** | $\frac{\text{Hinge\_res\_neg}}{\text{Res\_neg}} * 100$ |
| **Dipole moment/hyd moment ratio** | The ratio of dipole moment over the hydrophobic moment to describe the balance of polar versus nonpolar distributions per molecule. This was previously identified as an intrinsic non-redundant descriptor for a dataset of commercial mAbs. [viii] |
| **Ionic/hydrophobic patch area ratio** | The ratio of ionic patch area to hydrophobic patch area. This was previously identified as an intrinsic non-redundant descriptor for a dataset of commercial mAbs. [viii] |
| **BSA** | The buried surface area (BSA) of all chains in $Å^2$. This was calculated through the descriptors function in the BioMOE module in MOE 2020. Default parameters were used with no sampling. This was calculated off This was calculated at default values of pH 7.4, temperature of 300K and a salt concentration of 0.1M.[iii] |
| **BSA_HC** | The buried surface area (BSA) between the heavy and light chains in $Å^2$. This was calculated through the descriptors function in the BioMOE module in MOE 2020. Default parameters were used with no sampling. This was calculated off This was calculated at default values of pH 7.4, temperature of 300K and a salt concentration of 0.1M.[iii] |
| **BSA_LC_HC** | The buried surface area (BSA) between the heavy and light chains in $Å^2$. This was calculated through the descriptors function in the BioMOE module in MOE 2020. Default parameters were used with no sampling. This was calculated off This was calculated at default values of pH 7.4, temperature of 300K and a salt concentration of 0.1M.[iii] |

*Patch analysis of mAb 1 IgG1 and IgG3*

**Table S4 Analysis of positive, negative and hydrophobic surface patches for anti-IL8 IgG1 and IgG3 homology constructs.**

| Molecule | patch_hyd (Å$^2$) | patch_hyd% | patch_hyd_n | patch_pos % | patch_pos (Å$^2$) | patch_pos_n | patch_neg % | patch_neg (Å$^2$) | patch_neg_n | patch_ion (Å$^2$) | patch_ion_n | patch_cdr_hyd (Å$^2$) | patch_cdr_hyd_n | patch_cdr_pos (Å$^2$) | patch_cdr_pos_n | patch_cdr_neg (Å$^2$) | patch_cdr_neg_n | patch_cdr_ion (Å$^2$) | patch_cdr_ion_n | Asa_hyd (Å$^2$) | Res_ASA (Å$^2$) | Res_hyd (Å$^2$) | Res_pos (Å$^2$) | Res_neg (Å$^2$) |
|---|---|---|---|---|---|---|---|---|---|---|---|---|---|---|---|---|---|---|---|---|---|---|---|---|
| **IgG1** | 3790 | 42% | 65 | 33% | 2940 | 65 | 25% | 2250 | 40 | 5190 | 105 | 540 | 4 | 500 | 13 | 570 | 8 | 1070 | 21 | 33365 | 3747 | 57905 | 2924 | 2188 |
| **IgG3** | 5140 | 37% | 82 | 31% | 4240 | 81 | 32% | 4450 | 72 | 8690 | 153 | 560 | 4 | 460 | 12 | 530 | 9 | 990 | 21 | 40569 | 4450 | 70184 | 4232 | 4434 |

**Table S5 Analysis of positive, negative and hydrophobic surface patches for the hinge regions of anti-IL8 IgG1 and IgG3 homology constructs.**

| Molecule | Hinge_res_ASA (Å$^2$) | Hinge_res_hyd (Å$^2$) | Hinge_res_pos (Å$^2$) | Hinge_res_neg (Å$^2$) | Hinge_res_ASA contribution to res_ASA | Hinge_res_hyd contribution to res_hyd | Hinge_res_pos contribution to res_pos | Hinge_res_neg contribution to res_neg |
|---|---|---|---|---|---|---|---|---|
| **IgG1** | 958 | 50 | 35 | 77 | 2% | 1% | 1% | 4% |
| **IgG3** | 5902 | 456 | 375 | 493 | 8% | 9% | 9% | 11% |

*Physicochemical descriptors for anti-IL8 IgG1 and IgG3*

**Table S6 Sequence and structure based descriptors computed for anti-IL8 IgG1 and IgG3 homology constructs.**

| Molecule | pI_seq | pI_3D | Net Charge | Dipole Moment | Zeta Potential | Hydrophobicity Moment | Hydrophobic imbalance | Hydrophobic index | Dipole moment/ hyd moment ratio | Ionic/ hyd patch area ratio | BSA | BSA_HC | BSA_LC_HC |
|---|---|---|---|---|---|---|---|---|---|---|---|---|---|
| **IgG1** | 6.67 | 7.97 | 22.68 | 704.98 | 4353.5 | 15.26 | 0.71 | 1.00 | 0.16 | 1.37 | 5160.49 | 1636.04 | 3473.99 |
| **IgG3** | 6.54 | 7.84 | 21 | 2788.2 | 7340.2 | 6.21 | 1.08 | 0.94 | 0.38 | 1.69 | 5744.45 | 2030.33 | 3714.12 |

### Biophysical analysis of IgG1 and IgG3

#### *Analysis of identity by mass spectrometry*

The sequence and composition of anti-IL8 IgG1 and IgG3 was verified using peptide fingerprinting mass spectrometry. 250 µg of each molecule was denatured, reduced and alkylated. Further reduction and desalting to exchange samples into the digestion buffer followed, with trypsin digest (sequencing-grade, Promega, WI, USA) at a 1:20 (w/w) ratio after 2 hours of incubation. Mass spectrometry-liquid chromatography (MS-LC) was performed using an Orbitrap Exploris™ 240 Mass Spectrometer (Thermo Fisher Scientific, MA, USA), controlled by Xcalibur software (version 4.4.16.14, Thermo Fisher Scientific, MA, USA). A ACQUITY UPLC PEPTIDE CSH C18 (Waters, US) column was used for peptide separation, with 214nm absorbance detection. Separation of digested peptides was achieved by a method comprising of step wise gradients of buffer B, containing acetonitrile. The MS system was operated in positive ion mode, with a 200-2000 m/z scan range and a final resolution target of 15,000. Filtering criteria included charge states of 2-8 and minimum of 5 scans. Byos software (version 5.0-88 (2022.12), Protein Metrics, CA, USA) was used to process peptide fragments with precursor mass tolerance set at 20 ppm, fragment mass tolerance 1 and 2 set at 20 ppm and cleavage sites set as arginine and lysine. The post translation modifications (PTMs) screened for were methylation, oxidation, deamidation and pyroglutamate formation.

**Table S7 Verification of anti-IL8 IgG1 and IgG3 identity by peptide fragmentations.** Trypsin digest of this peptide following the same methodology showed coverage of this missing peptide, ensuring full identity verification. For post-translational modifications (PTMs), the % detection was relative to only peptides with expected full enzyme cleavage. PTMs with relative detection were noted. HC: Heavy chain; LC: Light chain; mwt: molecular weight; PTM: post-translational modification

| Molecule | LC coverage (%) | HC coverage (%) | LC mwt (Da) | HC mwt (Da) | LC PTMs | HC PTMs |
|---|---|---|---|---|---|---|
| IgG1 | 97.66 | 96.66 | 23433.83 | 49204.09 | M4 oxidation (0.4%) | M81 oxidation (0.2%), N317 deamidation (0.6%), M254 oxidation (4.2%), N363 deamidation (0.4%), M430 oxidation (1.8%) |
| IgG3 | 97.66 | 98.79 | 23433.83 | 54375.06 | M4 oxidation (0.6%), Q147 pyroglutamate (29.6%) | M81 oxidation (0.6%), M301 oxidation (6.9%), N263 deamidation (0.5%), N410 deamidation (0.5%), M446 oxidation (1.4%), N470 deamidation (0.2%), S473 methylation (1.7%), M477 oxidation (2.9%) |

*Scattering intensity profiles from nano-differential scanning fluorimetry (nano-DSF) of anti-IL8 IgG1 and IgG3*

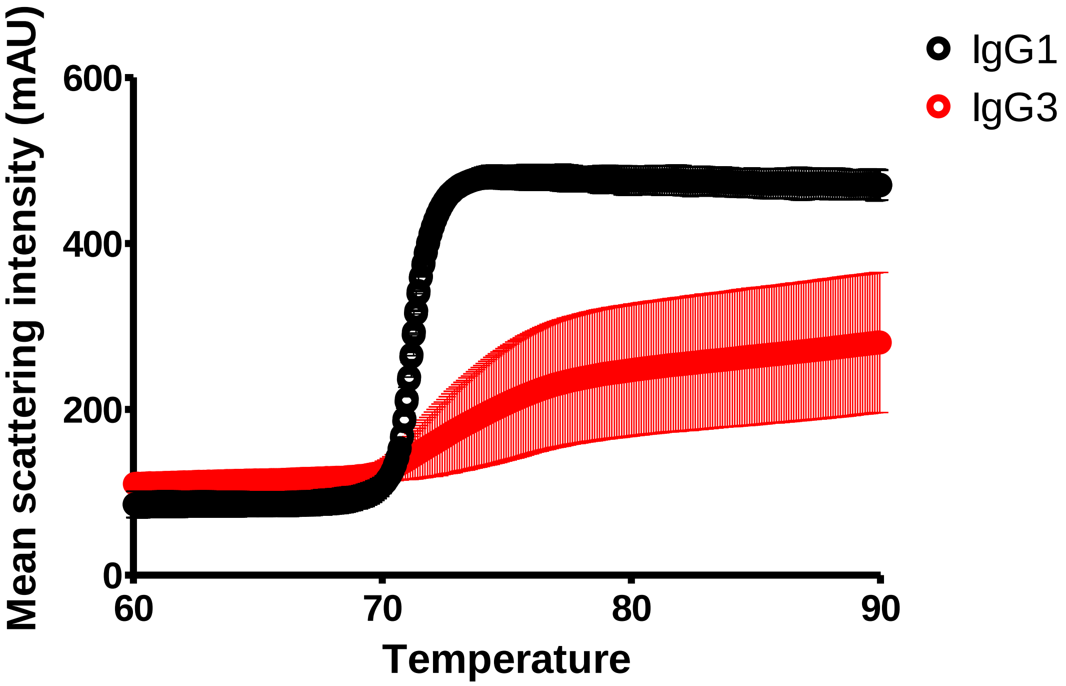

**Figure S1 Mean scattering intensity for anti-IL8 IgG1 (black) and IgG3 (red) differential scanning fluorimetry experiments**. IgG1 N=2, IgG3 N=3. Standard deviation error bars are shown.

*Diffusion coefficients from dynamic light scattering (DLS) for anti-IL8 IgG1 and IgG3*

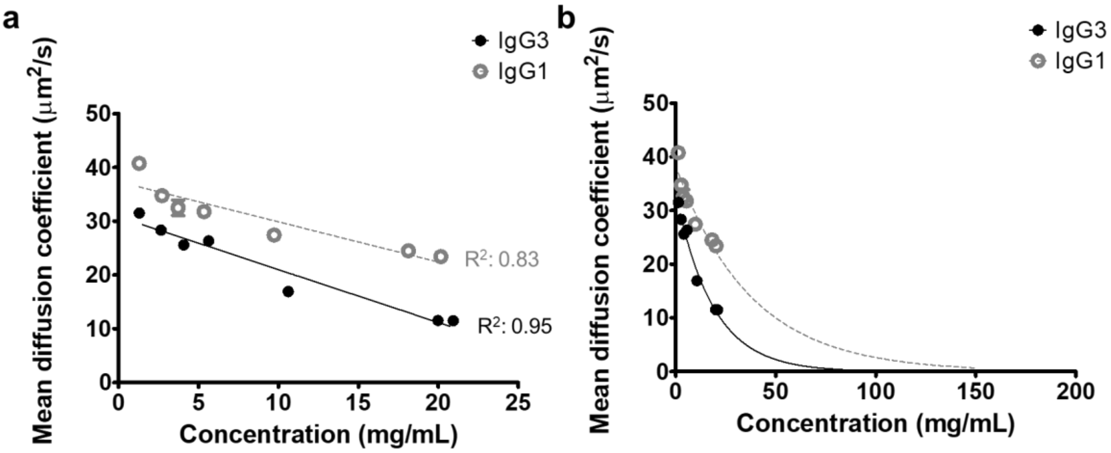

**Figure S2 Diffusion coefficients measured from dynamic light scattering (DLS) experiments**. **a,** linear regression of anti-IL8 IgG1 and IgG3 diffusion coefficients for self-interaction parameter, $k_D$, calculation. IgG1 linear equation: y = -746.89x + 37.36, IgG3 linear equation: y = -983.83x + 30.848. **b,** exponential growth fit of diffusion coefficients to extrapolate to 200mg/mL for theoretical viscosity calculations (Generalised Stokes Einstein equation). IgG1 exponential equation: y = 37.65e-0.025x, IgG3 exponential equation: y = 32.704e-0.052x.

***Intrinsic viscosity and the Huggins coefficient for anti-IL8 IgG1 and IgG3***

To determine the intrinsic viscosity, [η], and subsequently the Huggins coefficients for anti-IL8 IgG1 and IgG3 molecules, the relative viscosity, $\eta_{rel}$, was calculated (**Equation S1**).

$$\eta_{rel} = \eta/\eta_0 \qquad \textbf{(S1)}$$

Where the solution viscosity is $\eta_0$

Subsequently, the reduced viscosity, $\eta_{red}$, was calculated (**Equation S2)** and the intercept from the linear regression of $\eta_{red}$ over concentration (g/mL) determined [η] (**Figure S3)**. The Huggins coefficient was computed as described in **Equation 8**.

$$\eta_{red} = ((\eta_{rel} - 1)/c) \text{ x } 1000 \qquad \textbf{(S2)}$$

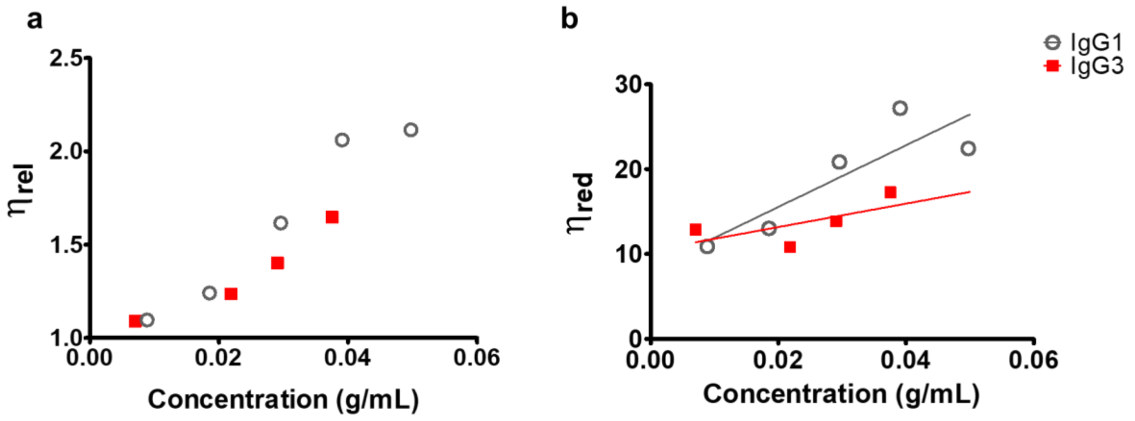

| Molecule | Slope | Intercept |
|---|---|---|
| IgG1 | 363.2 | 8.28 |
| IgG3 | 138 | 10.42 |

**Figure S3 a, Relative ($\eta_{rel}$) and b, reduced ($\eta_{red}$) viscosity for anti-IL8 IgG1 (grey) and IgG3 (red) over concentration (g/mL) in the dilute regime.** Linear regression for reduced viscosity over concentration determined the intrinsic viscosity (intercept).

[i] Chemical Computing Group ULC (2021). "MOE 2020.09: Ensemble Protein Properties."

[ii] Salgado, J. C., et al. (2006). "Predicting the behaviour of proteins in hydrophobic interaction chromatography: 1: Using the hydrophobic imbalance (HI) to describe their surface amino acid distribution." Journal of Chromatography A 1107(1): 110-119.

[iii] Long, W. (Chemical Computing Group) (2022). "Bio-MOE: Custom MOE Biologics Applications

[iv] Kyte, J. and R. F. Doolittle (1982). "A simple method for displaying the hydropathic character of a protein." Journal of Molecular Biology 157(1): 105-132.

[v] Sharma, V. K., et al. (2014). "In silico selection of therapeutic antibodies for development: viscosity, clearance, and chemical stability." Proc Natl Acad Sci U S A 111(52): 18601-18606.

[vi] Tanford, C. (1962). "Physical chemistry of macromolecules. , John Wiley & Sons, Inc., New York 16, N. Y., 1961." 51(2): 190-190.

[vii] Sillero, A. and J. M. Ribeiro (1989). "Isoelectric points of proteins: theoretical determination." Anal Biochem 179(2): 319-325.

[viii] Ahmed, L., et al. (2021). "Intrinsic physicochemical profile of marketed antibody-based biotherapeutics." Proceedings of the National Academy of Sciences 118(37): e2020577118.
